# Tripartite synergy - Metabolic crosstalk between two bacterial mutualists and a marine microalga promotes algal fitness

**DOI:** 10.64898/2026.05.29.728686

**Authors:** Bertille Burgunter-Delamare, Matthias Ostermeier, Trang Vuong, Patrick Then, Timm Yakin, Jörg Nickelsen, Mar Benavides, Maria Mittag

**Affiliations:** Matthias Schleiden Institute of Genetics, Bioinformatics and Molecular Botany, Friedrich Schiller University Jena, 07743 Jena, Germany; Molecular Plant Science, Ludwig-Maximilians-Universität München, 82152 Planegg-Martinsried, Germany; Cluster of Excellence Balance of the Microverse, Friedrich Schiller University, 07743 Jena, Germany; National Oceanography Centre, European Way, Southampton, SO14 3ZH, United Kingdom; Aix Marseille Univ., Université de Toulon, CNRS, IRD, MIO, Marseille, France

## Abstract

Marine microalgae form major parts of phytoplankton and are highly relevant for global CO_2_ fixation. Although microalgae have lived together with bacteria in the oceans for billions of years, these ecosystem-relevant interactions remain largely uncharacterized. Here, we have studied biotic interactions between two marine bacteria and a marine microalga. We show that an N_2_-fixing *Vibrio* provides ammonium for *Chlamydomonas* sp. and a *Marinobacterium.* Both microorganisms cannot survive in an ammonium-free environment. In exchange, the microalga promotes the growth of both bacteria, via secretion of heat-resistant metabolites in case of *Marinobacterium*. Reciprocally, the *Marinobacterium* releases heat-resistant metabolites that stimulate algal growth and increase its photosynthetic pigments, Photosystem II quantum yield, and starch accumulation. Electron microscopy reveals a strengthened starch sheath around the algal pyrenoid and indicates a modified periplasmic space for metabolic exchange. Our data highlight a tight synergy of a marine microbial trio promoting each other’s growth and algal fitness.

## Main

Photosynthetic protists, commonly known as microalgae, are major contributors to global CO_2_ fixation in the oceans and form the basis of marine food webs (Field et al., 1998). In their native ecosystems, microalgae have coevolved with diverse bacterial communities for billions of years, leading to intricate associations within the phycosphere, the nutrient-rich microenvironment surrounding the algal cell (Bell and Mitchell, 1972). These associations can modulate algal metabolism and growth in a commensal, mutualistic or antagonistic way (Burgunter-Delamare et al., 2024), thereby shaping aquatic ecosystems (Cirri and Pohnert, 2019). Overall, the activities of marine microorganisms including phytoplankton, bacteria, grazers and viruses involve marine microbial metabolites that can play a role in ecology and biogeochemistry (Moran et al., 2022). Gaining a detailed understanding of microbial interactions with many partners is challenging. Bottom-up or top-down approaches can be employed to explore these interactions. The first approach provides a mechanistic understanding of specific roles of microbial partners within defined cocultures. The second approach involves studying all relevant partners in a given environment, for example through studying annual patterns of plankton community succession (Deng et al., 2022) to understand the ecological relevance of these assemblages. In the latter, it may be hard to dissect the individual roles of a given partner. In contrast, the first approach can address the increasing complexity of algal-bacterial interactions in a step-wise manner. Conventionally, bottom-up studies have relied on bipartite models, using one alga and one bacterium. For instance, algal-bacterial mutualism has been shown with the freshwater soil alga *Chlamydomonas reinhardtii* (Calatrava et al., 2024) or the marine alga *Ostreococcus tauri* (Cooper et al., 2019) and a specific bacterium for each alga. By this way, we can gain important insights into the nature of the algal-bacterial interaction. However, bottom-up approaches do not provide detailed insights into the physical and chemical interactome of all microbial partners within a complex microbiome. Recently, tripartite studies have bridged the gap between bipartite systems and complex microbiomes, adding an additional layer of biological complexity and revealing previously hidden interaction modes. Thus, interactions among a cyanobacterium and viruses have been shown to result in metabolite release and elicit a chemotactic response in heterotrophic bacteria (Henshaw et al., 2024; Focardi et al., 2025). Likewise, tripartite interactions of the microalga *C. reinhardtii* and an antagonistic as well as a mutualistic bacterium can clarify algal survival strategies (Carrasco Flores et al., 2024). In addition, abiotic factors like nutrient availability and vitamins can play significant roles for microalgae and may influence algal-bacterial relationships (Croft et al., 2005; Harrison et al., 2025).

Here, we used a newly developed marine model *Chlamydomonas* species with a draft genome available (Carrasco Flores et al., 2021) to study algal-bacterial interactions. *Chlamydomonas* sp. was isolated in the coastal waters of Nantucket Sound (USA). *Chlamydomonas* sp. requires ammonium as a nitrogen source because it lacks genes encoding nitrate and nitrite reductases (Carrasco Flores et al., 2021). Therefore, we established a minimal artificial microbiome of the photoautotroph alga with a diazotroph and heterotrophic bacteria living in similar habitats (Duncan et al., 2022). We studied these microorganisms in bi- and tripartite cultures. Besides cyanobacteria, heterotrophic N_2_-fixers such as *Vibrio* can convert N_2_ gas into ammonium and are known to play key roles in marine N_2_-fixation (Zehr and Capone, 2020; Bonnet et al., 2022). The chosen bacterium, *Vibrio diazotrophicus*, can use various organic and inorganic nitrogen sources (Crétin et al., 2025). *V. diazotrophicus* and *Vibrio* spp. are generally widespread in the marine environment and have been characterised through metagenomes of the Arctic and Atlantic Oceans (Fu et al., 2020; Duncan et al., 2022; Crétin et al., 2025). We found that the presence of *V. diazotrophicus* enables the growth of *Chlamydomonas* sp. in an ammonium-free medium. *V. diazotrophicus* lives with *Chlamydomonas* sp. in a mutualistic manner, as diazotroph growth is reciprocally increased in algal coculture. As further bacterial partners, we have analysed four different *Marinobacterium* spp. in algal cocultures. These bacteria are also broadly present in the marine environment, and were also found in metagenomes of the Arctic and Atlantic Oceans (Duncan et al., 2022), as well as in coastal areas (Lpsn.dsmz.de/genus/marinobacterium). *Chlamydomonas* sp. lives in a symbiotic manner with all four *Marinobacterium* species. In each of these cocultures, *Marinobacterium* spp. enhanced algal growth and photosynthetic pigment concentrations. In return, the algal cells supported the growth of all four *Marinobacterium* spp.. In the case of *M. stanieri*, this mutualism is mediated by heat-resistant compound(s) secreted from both, the algal and bacterial partner, into the medium. The bacterial exometabolite(s) that are secreted from bacterial mono- and cocultures with the alga stimulate not only algal growth but also its photosynthetic capacity, including photosynthetic pigments, the quantum yield of Photosystem II, as well as starch production, as confirmed by electron microscopy (EM). EM also revealed that the exometabolite(s) induce an expansion of the periplasmic space in the microalgae that may facilitate metabolic exchange.

Bipartite studies with *V. diazotrophicus* and *M. stanieri* and the introduction of tripartite interactions of *Chlamydomonas* sp., *V. diazotrophicus* and *M. stanieri* revealed a tight synergy among all partners. Thus, *V. diazotrophicus* not only supports the growth of *Chlamydomonas* sp. in an ammonium-free environment but also the growth of *M. stanieri*. Notably, the combined presence of both bacteria further enhances algal growth and photosynthetic pigment concentration, establishing a highly interdependent minimal consortium of the three partners. These data highlight that the trio operates through sophisticated metabolic crosstalk, forming a robust functional unit that establishes a microbial “ménage-à-trois”. The trio ensures algal survival and fitness, as well as the survival of one bacterium and improved growth of both bacteria in an ammonium-free environment.

## Results

### A diazotrophic bacterium enables the growth of a marine ammonium-auxotroph alga while the alga supports bacterial growth in return

*Chlamydomonas* sp. lacks several genes involved in nitrate and nitrite metabolism (Carrasco Flores et al., 2021) (Fig. 1a). As a consequence, this alga depends on external ammonium as a nitrogen source (Fig. 1b, c; Extended Data Fig. 1). In coastal waters, primarily nitrate is often the dominant form of bioavailable nitrogen, as accessible ammonium is usually present at low or undetectable concentrations (Zehr and Kudela, 2011). The question arises as to how *Chlamydomonas* sp. survives without ammonium. One possibility would be that the alga obtains ammonium from diazotrophs, which are important contributors to nitrogen availability in coastal regions (Fulweiler et al., 2025). To test this hypothesis, we cocultured *Chlamydomonas* sp. with *V. diazotrophicus* in an ammonium-free medium. Impressively, *Chlamydomonas* sp. could grow in the presence of *V. diazotrophicus* to a similar extent as in ammonium-saturated medium (Fig. 1b, c; Extended Data Fig. 1; Methods), indicating that diazotrophy-derived nitrogen is enough to sustain the nitrogen needs for the alga.

**Fig. 1.**
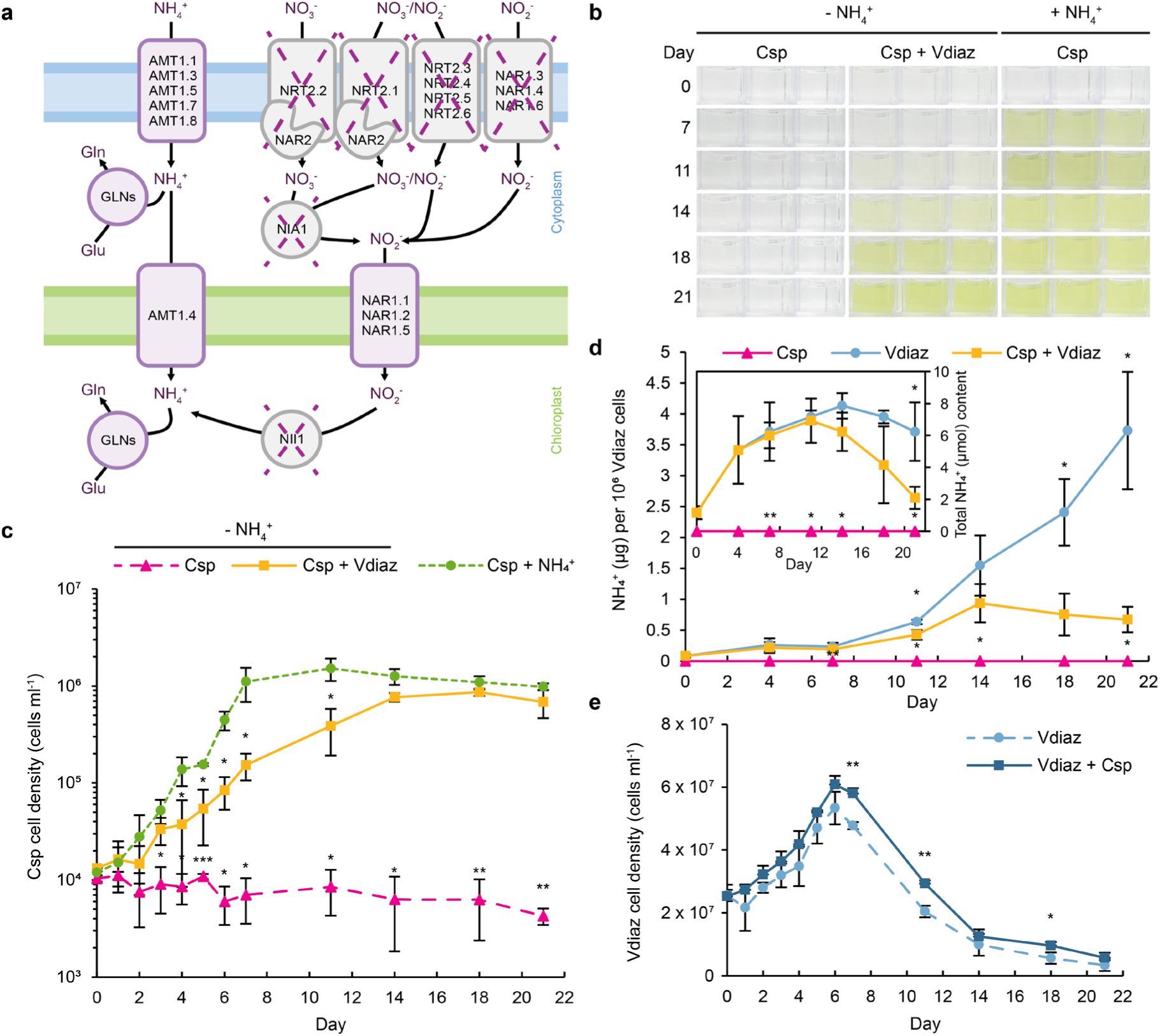
*V. diazotrophicus* (Vdiaz) enables the growth of *Chlamydomonas* sp. (Csp) by releasing NH_4_^+^ into the medium, while the alga supports bacterial growth in return. **a**, Schematic of relevant genes/proteins for green algal nitrogen metabolism, including transporters (Nrt, Nar, Amt) in the plasma- and chloroplast membranes, as well as nitrate reductase (Nia), nitrite reductase (Nii), and glutamine synthetases (Gln). Conserved gene sequences in *Chlamydomonas* sp. are shown in purple; missing ones are crossed out (dashed lines). Adapted from Carrasco Flores et al. (2021); functional aspects are based on *C. reinhardtii* (Schmollinger et al., 2014; Sanz-Luque et al., 2015). **b**, Algal mono- and cocultures. Organisms were cultivated in an NH_4_^+^-depleted YBCII medium over 21 days (Methods). An axenic culture of Csp in an NH_4_ -supplemented YBCII medium was used as a positive control. The full overview of the mono- and cocultures is shown in Extended Data Fig. 1. c, Algal cell densities in mono- and cocultures. For culturing, see **(b)**. Cells from a 10 ml cell suspension were harvested over 21 days. Genomic DNA was extracted and used for qPCR to determine cell densities (Methods). The shown statistics refer to Csp grown in the medium with ammonium (+NH_4_^+^). **d**, Normalised total NH_4_^+^ content per 10^6^ Vdiaz cells in mono- and cocultures over 21 days. Inlet: Total NH_4_^+^ content in mono- and cocultures over 21 days. The statistics refer to cocultures of Csp and Vdiaz. e, Bacterial cell densities in mono- and cocultures. For cultivation and determination of the cell densities, see the legends for (b) and (**c**). **b-e**, All experiments were done with *n* = 3 independent biological replicates. **c-e**, Error bars indicate SDs. Asterisks indicate significant differences as calculated by Student’s *t*-test: \*\*\**P* <0.001, \*\**P* <0.01, \**P* <0.05. Full statistical analyses are detailed in Extended Data Table 1a-c.

To characterise the exchange, we quantified extracellular ammonium levels in *V. diazotrophicus* mono- and cocultures, using an algal monoculture as a negative control. All cultures were grown in an ammonium-free medium (Fig. 1d). In the presence of the diazotroph bacterium, an increasing amount of ammonium was detected per 10^6^ bacterial cells. Conversely, extracellular ammonium levels were significantly lower in the coculture with *Chlamydomonas* sp. (Fig. 1d). This reduction indicates that the ammonium released by *V. diazotrophicus* is assimilated by *Chlamydomonas* sp..

We also examined the growth of *V. diazotrophicus* in mono- and algal cocultures (Fig. 1e). The bacteria reached their maximal growth on day 6-7 in both mono- and cocultures, providing 0.23 µg and 0.17 µg ammonium per 10^6^ bacterial cells, respectively. On day 21, a maximal ammonium content is reached (3.73 µg per 10^6^ bacterial cells in monoculture), but the bacterial cell density declined substantially, indicating that many bacteria have died. Nevertheless, even the smaller fraction of ammonium released between days 2 and 7 appeared to be sufficient to promote algal growth (Fig. 1c). The rapid decline of bacterial cell density after day 7 may be caused by the activation of a *V. diazotrophicus* prophage, resulting in bacterial lysis and a rapid release of ammonium (Mahoudeau et al., 2026). The maximum total ammonium content of the *V. diazotrophicus* monoculture is reached on day 14 and then declines again (Fig. 1d, inlet), suggesting a rapid lysis event after the growth peak. The algae clearly profit from this environment as the ammonium content gets significantly reduced in coculture until day 21 (Fig. 1d, inlet).

In return, the growth of *V. diazotrophicus* was slightly but significantly enhanced in coculture with *Chlamydomonas* sp. (Fig. 1e), supporting the idea of a mutualistic relationship between the two partners.

### Several species of the genus *Marinobacterium* establish mutualistic associations with *Chlamydomonas* sp. through reciprocal growth promotion and algal pigment enhancement

In its native ecosystem, *Chlamydomonas* sp. cohabitates with a variety of different bacteria, such as members of the genus *Marinobacterium* (Duncan et al., 2022). We investigated the influence of different *Marinobacterium* spp. available in microbial culture collections on *Chlamydomonas* sp. and included a marine *Pseudomonas* species, because *C. reinhardtii* is antagonised by *Pseudomonas* spp. (Aiyar et al., 2017; Carrasco Flores et al., 2024). While the marine *Pseudomonas deceptionensis* does not alter the growth of *Chlamydomonas* sp. significantly (Extended Data Fig. 2a, b), all examined *Marinobacterium* species enhance algal growth significantly and to a high extent. This is visible by the intensified green colour of the cultures (Fig. 2a; Extended Data Fig. 2a) and the strongly increased algal cell densities (Fig. 2b). The cooperative bacteria include *M. stanieri*, formerly known as *Pseudomonas stanieri* (Baumann et al., 1983; Satomi et al., 2002), which was first studied, as well as *M. litorale*, *M. mangrovicola* and *M. rhizophilum* (Methods).

**Fig. 2.**
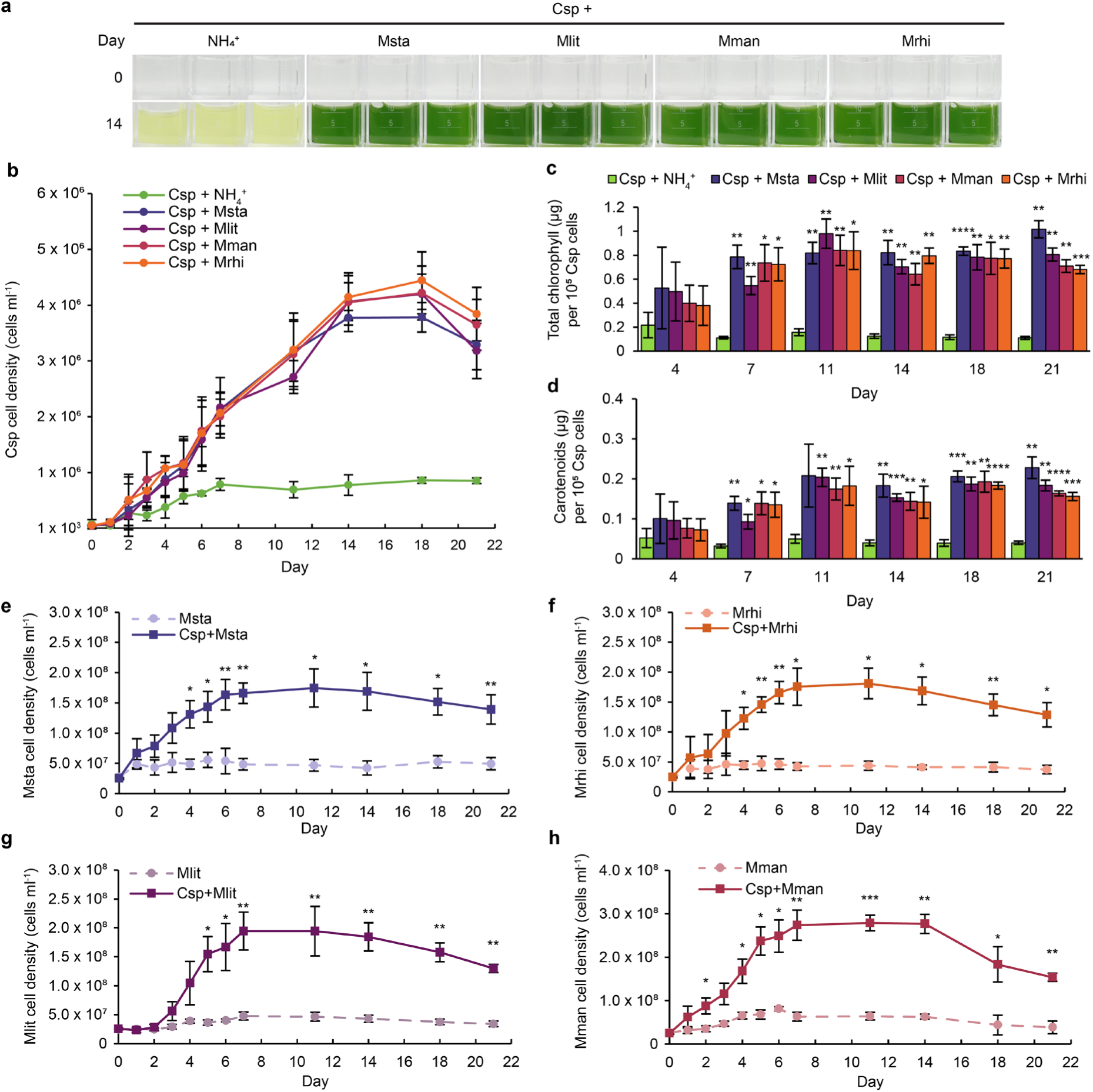
Several *Marinobacterium* spp. interact mutualistically with *Chlamydomonas* sp. (Csp). **a**, Algal mono- and cocultures with *Marinobacterium* spp. Organisms were cultivated in an NH_4_^+^-supplemented YBCII medium over 21 days (see Methods). An axenic culture of Csp in an NH_4_^+^-supplemented medium was used as a positive control. Msta: *M. stanieri*, Mlit: *M. litorale*, Mman: *M. mangrovicola*, Mrhi: *M. rhizophilum*. A comprehensive overview of all cocultures is shown in Extended Data Fig. 2a. **b**, Algal cell densities in mono- and cocultures. For cultivation, see legend of (**a**). Cells from a 10 ml cell suspension were harvested over 21 days. Genomic DNA was extracted and used for qPCR to determine cell densities (Methods). **c-d**, Normalised total chlorophyll (**c**) and carotenoids (**d**) contents per 10^5^ cells in Csp mono-and cocultures. The shown statistics refer to Csp grown in NH_4_^+^-YBCII. **e-h**, Bacterial cell densities of Msta (**e**), Mrhi (**f**), Mlit (**g**) and Mman (**h**) in mono- and cocultures. For cultivation, harvest and determination of cell densities, see legend of (**a**-**b**). **a-h**, All experiments were done with *n* = 3 independent biological replicates. **b-h**, Error bars indicate SDs. Asterisks indicate significant differences as calculated by Student’s *t*-test: \*\*\*\**P* <0.0001, \*\*\**P* <0.001, \*\**P* <0.01, \**P* <0.05. Full statistical analyses are detailed in Extended Data Table 1d-j.

We next asked whether the intensified green colouration reflected only higher cell numbers or also increased photosynthetic pigment production. Indeed, chlorophyll and carotenoid levels were significantly increased in cocultures compared to an algal monoculture from day 7 onwards. This was observed with all four *Marinobacterium* species (Fig. 2c, d), but not with *P. deceptionensis* (Extended Data Fig. 2c, d). Thus, all four *Marinobacterium* species not only support algal growth strongly but also enhance the algal photosynthetic pigments.

The support of algal growth in coculture raised the question whether the algal cells would also support bacterial growth, as would be the case in a mutualistic relationship. Indeed, *Chlamydomonas* sp. supports the growth of all four *Marinobacterium* species significantly and to a high level (Fig. 2e-h). Interestingly, the algal cells support the growth of *P. deceptionensis* significantly only on some days (Extended Data Fig. 2e).

### Heat-resistant exometabolites from bacteria in mono- and cocultures, as well as from algae in coculture, mediate a positive metabolic crosstalk among *Chlamydomonas* sp. and *M. stanieri*

In a next step, we studied the mechanism by which the bacteria may promote algal growth, taking *M. stanieri* as further example. One possibility is that they release metabolite(s) that the algae use to enhance their growth. To distinguish whether these metabolites are only produced in monoculture or also in coculture, we incubated the algae with spent media from bacterial mono- and cocultures. In all cases, we observed an intensification of the green colour as observed in the presence of *M. stanieri* (Fig. 3a, Extended Data Fig. 3a). Thus, the bacteria seem to release at least one metabolite that is needed to enhance algal growth and/or pigment production. Improved algal growth was corroborated by algal growth data with bacterial spent medium from mono- and cocultures (Fig. 3b, c). We also heat-treated the spent media (see Methods) to find out whether the exometabolite(s) are heat-stable. The addition of the heat-treated spent medium from bacterial mono- or cocultures caused the same positive algal growth effect (Fig. 3a-c), suggesting that the metabolite(s) released are heat-resistant.

**Fig. 3.**
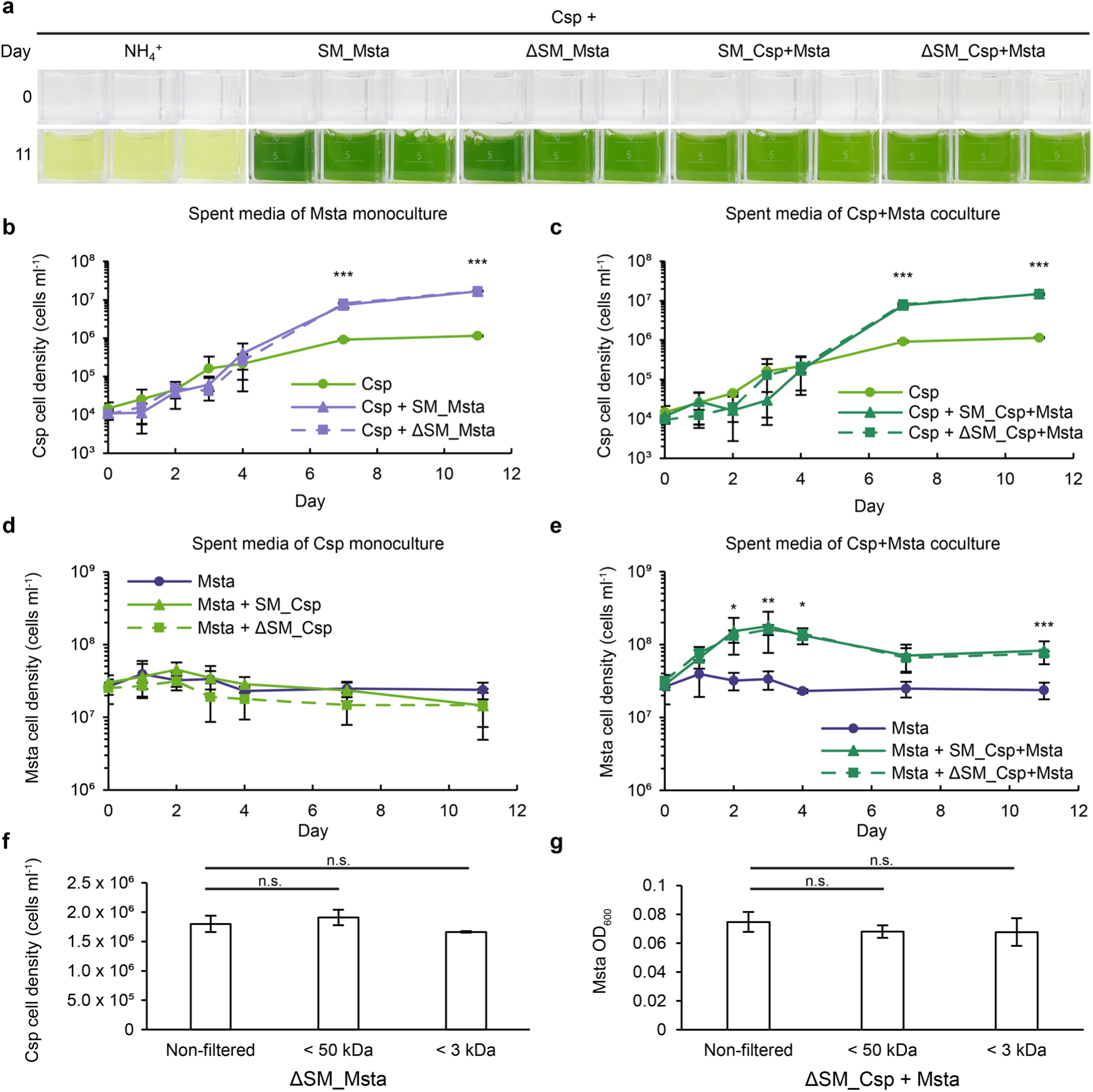
Small heat-resistant compounds are involved in the metabolic crosstalk of *M. stanieri* (Msta) and *Chlamydomonas* sp. (Csp). **a**, Algal monocultures in different spent media (SM). For a full overview, see Extended Data Fig. 3a. Csp was grown in bacterial spent media (SM_Msta) or in coculture spent media (SM_Csp+Msta) over 11 days. Heat-treated spent media (Methods) are indicated by “Δ” (**a-g**). Axenic cultures of Csp in NH_4_^+^-YBCII medium were used as a positive control (**a-e**). **b-c**, Algal cell densities of Csp grown in bacterial spent media (**b**) or coculture spent media (**c**), as described in (**a**), over 11 days. Cells from a 10 ml cell suspension were harvested over 11 days. Genomic DNA was extracted and used for qPCR to determine cell densities (Methods). Only the statistics of Csp grown in ΔSM_Msta (**b**) or in ΔSM_Csp+Msta (**c**), compared with Csp grown in NH_4_^+^-YBCII, are shown. **d-e**, Bacterial cell densities. Msta was grown in algal monoculture spent media (SM_Csp; **d**) or coculture spent media (SM_Csp+Msta; **e**) over 11 days. Only the statistics of Msta grown in ΔSM_Csp (**d**) or in ΔSM_Csp+Msta (**e**), compared with Msta grown in NH_4_^+^-YBCII, are shown. **f**, Algal cell densities after one week in non-filtered heated bacterial spent media (ΔSM_Msta) or after size fractionation (< 50 or < 3 kDa). The shown statistics are compared with Csp grown in non-filtered ΔSM_Msta. **g**, Bacterial cell densities after one week in different fractions of ΔSM_Csp+Msta (see **f**). The shown statistics are compared with Msta grown in non-filtered ΔSM_Csp+Msta. **a-g**, All experiments were done with *n* = 3 independent biological replicates. **b-g**, Error bars indicate SDs. Asterisks indicate significant differences as calculated by Student’s t-test: \*\*\**P* <0.001, \*\**P* <0.01, \**P* <0.05; n.s., not significant. Full statistical analyses are detailed in Extended Data Table 1k-n.

Similarly, we examined whether the algae secrete metabolite(s) in mono- and/or coculture supporting bacterial growth. The growth promotion of *M. stanieri* could not be observed in spent media from algal monocultures (Fig. 3d). Only, compound(s) of spent medium from cocultures resulted in an increase of bacterial growth (Fig. 3e). This was also the case upon heat-treatment of the compounds. The cocultivation of the bacteria with the algae thus triggers the production of compound(s), which may present a mutual “bonus” for the positive effects sparked by the bacteria.

Size fractionation restricted the yet unknown product(s) from mono- and cocultures to a size of less than 3 kDa, with both the algal and bacterial compound(s). The bacterial compound(s) < 3 kDa were found in monoculture (Fig. 3f) and coculture (Extended Data Fig. 3b) along with their effects on the algae. < 3 kDa compound(s) coming from the algae were found in monoculture (Extended Data Fig. 3c) and coculture (Fig. 3g), but had only an effect on the bacteria from coculture, as mentioned before. As the secreted compounds from bacterial and algal cells are heat-resistant and smaller than 3 kDa, it is unlikely that a protein is involved. The results are rather indicative of a heat-resistant metabolite(s), possibly specialised natural product(s).

### Heat-resistant bacterial exometabolite(s) enhance photosynthetic efficiency, induce differential starch/carbon partitioning and extend the cell wall region

We next examined whether the heat-resistant secreted metabolite(s) of *M. stanieri* that elicit the algal green colour intensification (Fig. 3a) also increase the levels of the photosynthetic pigments in the algal cells. In the presence of metabolites from the spent bacterial medium (mono- and cocultures), both chlorophyll (Fig. 4a, Extended Data Fig. 4a) and carotenoid concentrations (Fig. 4b, Extended Data Fig. 4b) increased significantly on representative days 7 and 11 of growth (Extended Data Fig. 3a). This effect persisted following heat treatment of the spent medium, confirming the thermal stability of the active compound. The maximum quantum efficiency of Photosystem II (PSII), represented by the Fv/Fm value, was also enhanced upon the addition of heated bacterial spent medium of the bacterial monoculture on day 7 of growth (Fig. 4c). Additionally, starch levels increased more than twofold (Fig. 4d) as a result of the enhanced photosynthetic capacities.

**Fig. 4.**
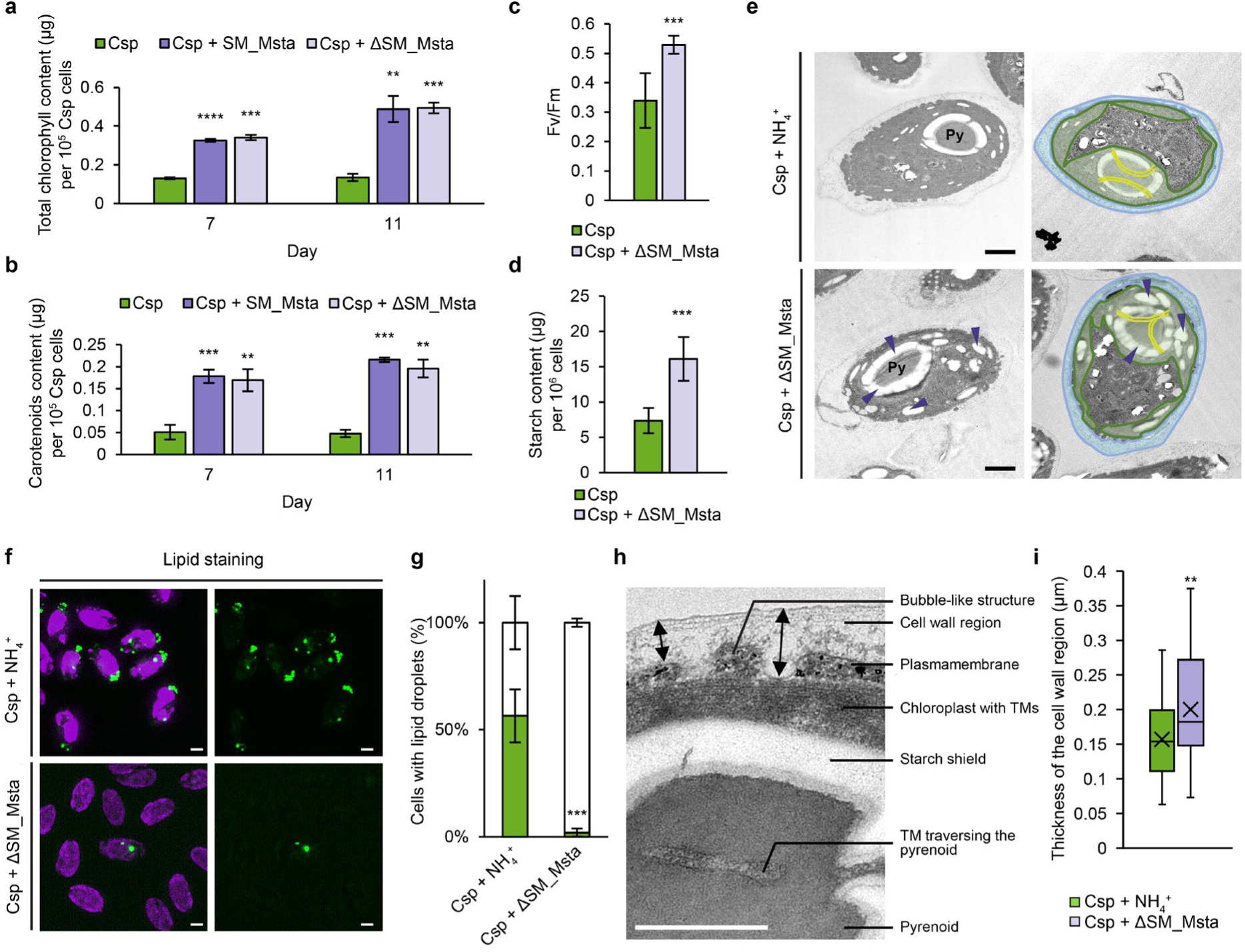
Heat-resistant bacterial product(s) enhance algal photosynthetic capacities and the thickness of the cell wall region. **a-b**, Normalised total chlorophyll (**a**) and carotenoids (**b**) content per 10^5^ cells in *Chlamydomonas* sp. (Csp) cells grown in NH_4_^+^-YBCII (**a-i**), in spent bacterial media (SM_Msta; **a-b**) or in heated spent media (ΔSM_Msta; **a-i**). The statistics refer to Csp grown in NH_4_^+^-YBCII. **c**, Maximum photochemical efficiency (Fv/Fm) of PS II of algal cells. Measurements were done in liquid cultures after normalising all cultures to 5 μg ml^-1^ *Chl*. **d**, Starch content of Csp cells. **e**, EM graphs of high-pressure frozen ultra-thin sections. On the right, the chloroplast is highlighted in green, with traversing thylakoid membranes (TMs) through the pyrenoid (Py on the left) and the cell wall area (light blue). Starch granules are indicated with purple arrows. Scale bars = 1 µm. **f**, Stained lipid droplets (green colour, right panels) and combined purple coloured *Chl* autofluorescence (left panels). SIM-Apotome images were taken by a ZEISS Elyra 7. **g**, Lipid droplet accumulation in Csp cells. **h**, EM picture of Csp+NH_4_^+^ with a detailed view of the different layers and a TM traversing the pyrenoid. The double-sided black arrows in the upper part illustrate the measurement shown in (**i**). Scale bar = 0.5 µm. **i**, Comparison of the thickness of the cell wall region. 43 random measurement points were taken per strain, each in 20 cells. Boxplots: centre line, median; cross marker, mean, box limits, upper and lower quartiles; whiskers, minimum and maximum values of the data set. **a-i**, Experiments were done with *n* = 3 (a-d, f-g) or *n* = 2 (e, h-i) independent biological replicates. **a-d, g, i**, Error bars indicate SDs. Asterisks indicate significant differences as calculated by Student’s *t*-test: \*\*\*\**P* <0.0001, \*\*\**P* <0.001, \*\**P* <0.01. Full statistical analyses are detailed in Extended Data Table 1o-p.

We also analysed the subcellular structures of *Chlamydomonas* sp. by EM on day 7 of growth in the absence and presence of the heated bacterial metabolite(s). *Chlamydomonas* sp. has an oval shape and a prominent pyrenoid surrounded by a starch sheath (Fig. 4e, upper left part). To highlight the plastid area, it was visualised by green colour (Fig. 4e, upper right part). In contrast to *C. reinhardtii*, which has a U-shaped chloroplast (Sasso et al., 2018), the plastid of *Chlamydomonas* sp. surrounds the entire oval algal cell directly beneath the cell wall area. Embedded within the chloroplast is the pyrenoid subtended by thylakoid membranes, which in turn shows no visible difference in any condition. Importantly, the starch sheath is strengthened in the presence of the heated bacterial metabolite(s) and further starch corns are present within the plastid, as indicated by arrows (Fig. 4e, lower parts, Extended Data Fig. 4c). As algal cells often prefer one type of energy storage compounds and may decrease others, we analysed the levels of lipid droplets by structured illumination microscopy (SIM) images in apotome mode (Fig. 4f, Extended Data Fig. 4d). Indeed, lipid droplets are significantly reduced in cells incubated with the heated bacterial metabolite(s) (Fig. 4g).

The cell wall region of *C. reinhardtii* includes the periplasmic space that is situated outside of the plasmalemma (Shimamura et al., 2024). The marine algal cell region seems sophisticated: In many cells, bubble-like structures can be recognised within the cell wall region extending from the cytoplasm into the periplasmic space. Although it is not yet clear whether these bubble-like structures that are surrounded by the plasma membrane are altered upon addition of spent medium, they seem to cause an extended surface area of the cytoplasm (Fig 4h). Of note is that the periplasmic space in the cell wall region is overall thickened in the presence of the heated bacterial metabolite(s) (Fig. 4i).

### A synthetic tripartite consortium enables algal cells to grow in an ammonium-free environment with enhanced algal pigment content

The metabolites of *V. diazotrophicus* and *M. stanieri* released in algal coculture enable algae to select habitats without ammonium and to strengthen algal photosynthetic capacity. To understand the interplay of all three microorganisms, we created a minimal artificial consortium, consisting of the two above-mentioned bacteria and *Chlamydomonas* sp..

The growth of all three microorganisms in a tripartite culture without ammonium was compared to the growth of *Chlamydomonas* sp. and *M. stanieri* cocultures with and without ammonium, respectively, as well as with a coculture of *Chlamydomonas* sp. and *V. diazotrophicus* without ammonium (Fig. 5a, Extended Data Fig. 5a). Indicative of the green colour of the algal cells, we observed a positive effect on algal growth in the tripartite culture with *M. stanieri* compared to a coculture of *Chlamydomonas* sp. and *V. diazotrophicus* alone (Fig. 5a). Algal growth curves exhibited significantly higher algal cell densities in the tripartite consortium (Fig. 5b).

**Fig. 5.**
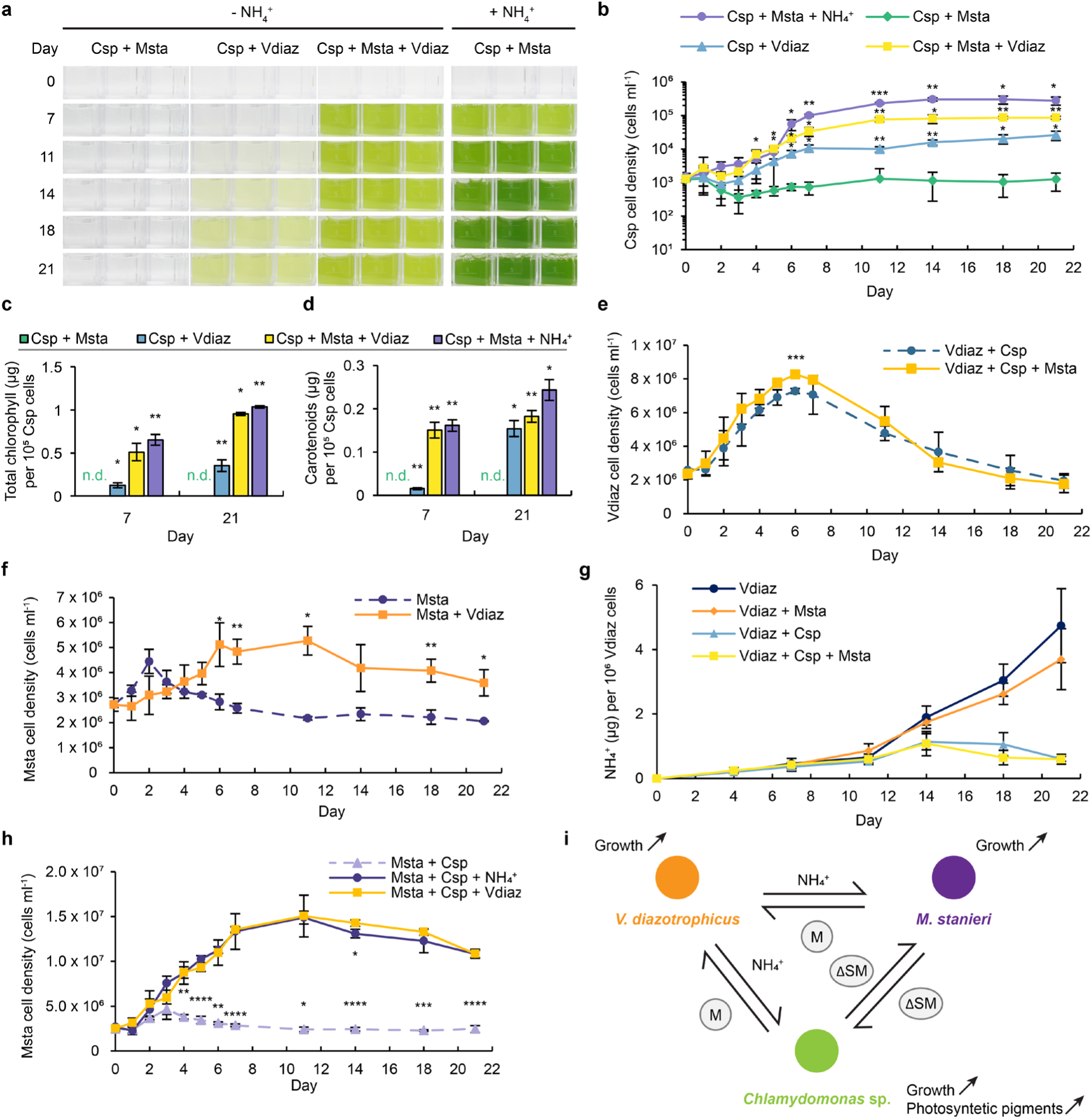
Strong synergistic effects are present among all three partners in a microbial “ménage-à-trois”. **a**, Algal mono- and cocultures. The organisms were cultivated in NH_4_^+^-depleted YBCII medium over 21 days (see Methods), unless otherwise indicated by (+ NH_4_^+^). Csp+Msta (*Chlamydomonas* sp. and *M. stanieri*) cocultures in an NH_4_^+^-supplemented YBCII medium were used as a positive control. The full overview of the mono, bi- and tripartite cultures is shown in Extended Data Fig. 5a. **b**, Algal cell densities in bi- and tripartite cocultures. For cultivation, see legend of (**a**). Cells from a 10 ml cell suspension were harvested over 21 days. Genomic DNA was extracted and used for qPCR to determine cell densities (Methods). **c-d**, Normalised total chlorophyll (**c**) and carotenoids (**d**) content per 10^5^ cells in Csp cocultures from days 7 and 21. n.d.: not detected. **e**, Cell densities of *V. diazotrophicus* (Vdiaz) in cocultures with Csp or with Csp and Msta. **e-h**, For cultivation, harvest and cell densities, see legends of (**a-b**). **f,** Cell densities of Msta in mono- and cocultures with Vdiaz. **g**, Normalised total NH_4_^+^ content per 10^6^ Vdiaz cells in mono- and cocultures, over 21 days. **h**, Bacterial cell densities of Msta in cocultures with Csp and in tripartite cultures with Csp and Vdiaz. **i**, Schematic of the Csp-Msta-Vdiaz tripartite interaction. Vdiaz provides NH_4_^+^ to Csp and Msta, allowing their growth in an NH_4_^+^-depleted medium. Msta and Csp support Vdiaz by providing unknown metabolite(s) (M). Msta enhances Csp growth and photosynthetic pigments by providing heat-resistant secreted metabolite(s) (ΔSM). Msta benefits from unknown ΔSM in cocultures with Csp. **a-f**, All experiments were done with *n* = 3 independent biological replicates. **b-h**, Error bars indicate SDs. Asterisks indicate significant differences as calculated by Student’s *t*-test: \*\*\*\**P* <0.0001, \*\*\**P* <0.001, \*\**P* <0.01, \**P* <0.05. Full statistical analyses are detailed in Extended Data Table 1q-w.

In the absence of ammonium, *Chlamydomonas* sp. showed no significant growth in the coculture with *M. stanieri*, while the addition of *V. diazotrophicus* (bipartite culture) supports algal growth significantly (Fig. 5a, b). In the presence of both bacteria (tripartite culture), algal growth is even further enhanced, as mentioned above. Yet, algal growth is not as high as within a bipartite culture with *M. stanieri* in the presence of ammonium, which has, however, a saturating NH ^+^ concentration of 17 mM. Importantly, the presence of *M. stanieri* in the tripartite culture ensures that the algal cells have enhanced photosynthetic pigments over the entire growth period (Fig. 5c, d; Extended data Fig. 5b, c).

The growth of *V. diazotrophicus* was further augmented in the tripartite culture relative to its bipartite coculture with *Chlamydomonas* sp. (Fig. 5e), while it was not augmented in a bipartite culture with *M. stanieri* (Extended Data Fig. 5d). Specifically, the presence of *M. stanieri* in the tripartite culture increased the peak cell density of *V. diazotrophicus* on day 6. However, this enhancement was transient, as bacterial cell densities in both the bi- and tripartite conditions converged during the subsequent decline phase (Fig. 5e).

We also analysed whether the other bacterial partner, *M. stanieri,* may profit from *V. diazotrophicus* when ammonium is missing. *M. stanieri* clearly profited from the presence of *V. diazotrophicus* (Fig. 5f). Obviously, *M. stanieri* fails to maintain robust growth in the absence of exogenous ammonium; there was only a marginal increase in cell density observed until day 2. This may be attributed to the utilisation of endogenous nitrogen reserves carried over from the precultures. Following day 2, however, the cell density of the *M. stanieri* monoculture declined and remained stagnant. In contrast, *M. stanieri* achieved significantly higher cell densities when cocultured with *V. diazotrophicus*. This indicates that the diazotroph supports *M. stanieri* growth through ammonium exudation. Consequently, extracellular ammonium abundance was lower in *M. stanieri* - *V. diazotrophicus* cocultures compared to monocultures. (Fig. 5g). The lowest ammonium concentration was observed in a bipartite culture with *V. diazotrophicus* and *Chlamydomonas* sp. as well as in a tripartite culture with *M. stanieri* in addition (Fig. 5g). These results provide further evidence that algal cells are the primary assimilators of the ammonium exuded from *V. diazotrophicus*. Nevertheless, *M. stanieri* growth was also significantly supported with ammonium within the tripartite culture (Fig. 5h). Summa summarum, the algal cells, as well as the bacteria, profit from each other through metabolic crosstalk, highlighting a strong mutual synergy in the tripartite interaction (Fig. 5i).

## Discussion

In this study, we demonstrate the mutual dependence of microbial partners in a nutrient-deficient environment lacking ammonium. Two of the partners, the microalga *Chlamydomonas* sp. and the heterotrophic bacterium *M. stanieri*, are ammonium auxotrophs and rely on diazotrophic bacterial partners such as *V. diazotrophicus* for survival. Reciprocally, the alga enhances the growth of the diazotrophic bacterium. This partnership is ecologically significant given that dinitrogen is one major form of nitrogen in the oceans, where nitrogen compounds are usually only available at low rates (Moore et al., 2013). Dinitrogen fixation by diazotrophs such as *V. diazotrophicus* is metabolically expensive, requiring 16 ATP molecules for the generation of a single molecule of ammonium, and consequently necessitates a robust genetic regulatory system (Dixon and Kahn, 2004; Crétin et al., 2025).

Our data suggest that *V. diazotrophicus* secretes only a fraction of this metabolically expensive nutrient to its partners during active growth (Fig. 1d-e). This amount is, however, sufficient to promote early growth of the algae and *M. stanieri* (Fig. 1c and 5f). The highest ammonium levels are observed during a rapid population decline phase, likely as a result of cell death. The fast decline in cell density of *V. diazotrophicus* could be related to the activation of a prophage of *V. diazotrophicus* (Mahoudeau et al., 2026). The activation of the prophage was observed in exponential and stationary phases. Thus, an ammonium-rich microenvironment is suddenly created (Fig. 1d, inlet), from which both the algae and the other bacterial partner profit to ensure best survival in an ammonium-free environment (Fig. 5f, g). Interestingly, also *V. diazotrophicus* profits from the presence of *M. stanieri* in the tripartite culture (Fig. 5e). This effect, however, is restricted to the peak cell density of *V. diazotrophicus*, indicating that certain metabolite(s) may be needed in a high concentration to promote the growth of *V. diazotrophicus*.

Besides ammonium, other secreted metabolites that are < 3 kDa and heat-resistant play important roles. *M. stanieri* bacteria release the metabolites constitutively, as they are found in bacterial mono- and cocultures (Fig. 3). In contrast, algal cells secrete the small heat-resistant metabolite(s) that promote bacterial growth of *M. stanieri* only in coculture (Fig. 2e). They may sense the bacterial growth-promoting effect and support it by secreting their own growth-promoting factors. We do not know yet the nature of the released metabolite(s) from the algae and bacteria. B vitamins can be exchanged in mutualistic algal-bacterial partnerships (Croft et al., 2005; Carrasco Flores et al., 2024), but they seem unlikely here, as they are light and/or heat-sensitive (Schnellbaecher et al., 2019; Herrera-Ardila et al., 2022) and some are produced in green algae (Carrasco Flores et al., 2024). Interestingly, the algae support bacterial species widely in cocultures, as growth-promoting effects for several *Marinobacterium* spp., as well as for *P. deceptionensis* were observed (Fig. 3, Extended Data Fig. 2), pointing to a bacterial-related metabolite.

Bacterial support of *Chlamydomonas* sp. in both bi- and tripartite cultures influences both algal biomass accumulation and the enhancement of photosynthetic pigments. Experiments using the spent medium of *M. stanieri* further demonstrate that bacterial exometabolites stimulate overall algal photosynthetic capacity (Fig. 4a-e). Besides photosynthetic pigments, the PSII quantum yield and starch content are strongly enhanced. EM revealed that the starch sheath around the pyrenoid is strengthened. In *C. reinhardtii*, the carbon concentrating mechanism (CCM) is closely associated with this starch sheath (Mackinder et al., 2017). A thicker starch layer is thought to maintain the carboxylase function of the RUBISCO by preventing CO_2_ leakage from the pyrenoid (Fei et al., 2022). Such a correlation may also be relevant for *Chlamydomonas* sp. The pyrenoid of this alga is very distinct (Fig. 4e) and situated on one extended side of the chloroplast. In contrast to *C. reinhardtii* (Sasso et al., 2018), the chloroplast of *Chlamydomonas* sp. is not U-shaped, but oval-shaped. Another intriguing feature is the broadened cell wall area in the presence of the bacterial spent medium. The bubble-like regions extending from the cytoplasm into the broadened periplasmic space of the cell wall area may play a role in metabolite exchange. The expansion of the cytoplasmic surface area may facilitate more efficient metabolite transfer, as in an arbuscular mycorrhiza, where branched fungal hyphae are involved (Delaux and Gutjahr, 2024). The broadened periplasmic space may also support metabolite transfer. In *C. reinhardtii*, the periplasm stores enzymes, like carbonic anhydrase 1 that is involved in the CCM mechanism (Shimamura et al., 2024), arylsulfatase in response to sulfur deprivation (De Hostos et al., 1989) or FEA1 (Fe assimilation) promoting iron delivery (Allen et al., 2007). The expanded cell wall area, including the periplasm, may also work as a special kind of phycosphere or biofilm reservoir for the bacteria to facilitate metabolic exchange. Another possibility is a potential conversion of this area to an extracellular matrix that is, for example, found in multicellular green algae like *Volvox* (Kirk, 2005). Extracellular matrix can attract heterotrophic bacteria by supplying nutrients and protecting them, as observed in the Jekyll-and-Hyde partnership between the haptophyte alga *Emiliana huxleyi* and *Phaeobacter inhibens* (Seyedsayamdost et al., 2011; Lipsman et al., 2024).

The artificial minimal microbiome established in this study via bi- and tripartite algal-bacterial interactions provides unique insights into the complex interplay of these microbial interactions in marine environments. It highlights that microalgae not only may rely on the support of bacteria and vice versa, but also reveals that the bacterial partners can profit from one another. These mutualistic microbial partnerships are further strengthened in a tripartite culture. Moreover, it becomes obvious that the survival of microorganisms in environments lacking organic nitrogen compounds strongly benefits from diazotrophic bacteria. In contrast to cyanobacteria, they do not have to compete with microalgal cells for carbon dioxide (Ji et al., 2017) and may thus be favoured in such consortia. The observed mutualisms are manifold and exemplify the importance of each of the three partners for the metabolic crosstalk that enables better growth and thus survival for all of them.

## Methods

### Algal strain and growth media

Experiments were carried out with *Chlamydomonas* sp. SAG 25.89, obtained from the Culture Collection of Algae at Göttingen University (SAG). It should be noted that *Chlamydomonas* sp. SAG 25.89 corresponds to strain CCMP235 (Aiyar et al., 2017; Carrasco Flores et al., 2021), isolated from Falmouth Great Pond (MA, USA) of Nantucket Sound (https://ncma.bigelow.org/CCMP235). The algal cells were grown in NH_4_ ^+^-YBCII medium (adapted from Chen et al. (1996) by adding 17 mM NH_4_Cl and 20 mM HEPES (Carrasco Flores et al., 2021). The pH of the medium was 6.65 to mimic brackish environmental conditions (Shortelle and Colburn, 1987). As indicated, the ammonium source was omitted in some cases.

### Bacterial strains and growth media

All bacteria were obtained from bacterial strain collections and grown initially in the indicated medium and at the advised temperature provided by the strain collection. For comparative studies of the main cultures, all bacteria were grown in the modified NH_4_ ^+^-YBCII medium, in the presence or absence of ammonium as further described under cultures below.

The bacterium *Vibrio diazotrophicus* (DSMZ 2605) was obtained from the DSMZ (Braunschweig, Germany). Axenic bacteria were precultured in Vibrio medium (VM, Medium 308 DSMZ; https://www.dsmz.de/microorganisms/medium/pdf/DSMZ_Medium308.pdf) at 26 °C with constant orbital shaking (200 rpm). The bacteria *Marinobacterium stanieri* (DSMZ 7027), *M. litorale* (DSMZ 23545), *M. mangrovicola* (DSMZ 27697) and *M. rhizophilum* (DSMZ 18822) were obtained from the DSMZ (Braunschweig, Germany). Axenic bacteria were precultured in Marine Broth (MB, DIFCO) at 26 °C with constant orbital shaking (200 rpm). The bacterium *Pseudomonas deceptionensis* (DSMZ 26521) was also obtained from the DSMZ. Axenic bacteria were precultured in Tryptic Soy Broth (TSB, Millipore) at 26 °C with constant orbital shaking (200 rpm).

### Correlation between OD_600_ and bacterial cell density

Bacteria were cultivated in the above-mentioned media (VM, TSB or MB) for precultures. OD_600_ was measured with a Jenway 6305 spectrophotometer using diluted bacterial cell suspensions. A series of dilutions was prepared. From each dilution, 200 μL were transferred to a VD, TSB or MB plate and incubated at 26 °C for three days until colonies were properly grown and visible for counting. The colony forming units (CFU) counts were then used with the OD_600_ to generate a calibration curve (Extended Data Fig. 6a).

### Mono-, bi- and tripartite cultures

*Chlamydomonas* sp. cells were precultured to a cell density of 3-6 x 10^5^ cells ml^-1^ for the mono-and cocultures. Cell counts were determined using a Countess 3 FL Automated Cell Counter (Thermo Fisher Scientific Inc.) and were verified and adjusted by an improved Neubauer chamber (no. T729.1; Carl Roth, Karlsruhe, Germany). *Chlamydomonas* sp. was grown in YBCII medium with or without NH_4_ ^+^ at 23 °C under a 12:12 light-dark cycle using white light (Osram L36W/840, lumilux, cool white, Osram) with a light intensity of about 60 μmol m^-2^ s^-1^ and orbital shaking (100 rpm). Bacterial precultures of *V. diazotrophicus, Marinobacterium* spp. and *P. deceptionensis* were grown in VD, MB and TSB medium, respectively. OD values were correlated with bacterial cell densities. Prior to cultivation, bacterial and algal cells were washed twice and resuspended in YBCII media with or without NH_4_ ^+^, as indicated, at a starting algal cell density of 1 x 10^4^ cells ml^-1^ and a bacterial cell density of 2.5 x 10^7^ cells ml^-1^ per bacterium, as bacteria can reach high cell densities in coastal areas. These cultures were grown under the conditions described above for *Chlamydomonas* sp. growth and photo-documented on a regular basis during the growth period.

### Spent Media

*Chlamydomonas* sp. and *M. stanieri* mono- and cocultures were prepared as mentioned above. At day 7, the cultures were centrifuged (20 min, 3500 x *g*) and the supernatant filtered (0.22 µm). The filtrate was used as spent medium; one batch was heat-treated (autoclaved at 121 °C for 35 min). Spent media from mono- and cocultures were used as cultivating media for the growth of *Chlamydomonas* sp. and *M. stanieri*, respectively. The cultures were grown for 11 days under the conditions described above for *Chlamydomonas* sp. growth. Cultures in fresh media were used as a control. The cultures were photo-documented on a regular basis during the growth period.

### DNA extraction and DNA-based cell quantification via qPCR

DNA extraction and cell quantification by qPCR were done as described in Carrasco Flores et al., (2024). Algal and bacterial samples were harvested by centrifugation at 3500 x *g* for 20 min. The supernatant was discarded, and the cell pellet was stored at -20 °C until extraction. For DNA extraction, the extraction buffer consisted of 100 mM Tris, 10 mM Na_2_EDTA, and 10x Yellow Sample Buffer, which was obtained 40x from ThermoFisher Scientific (No. R1381). The pH was adjusted to 8. An aliquot of 200 µL of extraction buffer was added to the frozen cell pellets. The pellets were thawed at room temperature for 30-40 min, thoroughly vortexed to full resuspension and incubated in a water bath at 100 °C for 10 min. The mixture was briefly vortexed and centrifuged at 3440 x *g*, 4 °C for 20 min. The supernatant was transferred to new vials and stored at -20 °C for further use, and the remaining cell debris was discarded.

Cell densities were determined by correlating the quantified DNA to its equivalent cell density using calibration curves for each microbial culture. Algal cultures were inoculated at a cell density of 4-6 x 10^5^ cells ml^-1^ and incubated for 3 to 4 days, reaching a density of about 1.0-1.2 x 10^6^ cells ml^-1^. Algal cultures were then adjusted to 10^5^ cells ml^-1^. The suspension was then serially diluted in NH_4_ ^+^-YBCII, resulting in cell densities of 10^4^, 10^3^, and 10^2^ cells ml^-1^. Liquid cultures of *M. stanieri* and *P. deceptionensis* were adjusted to 10^8^ cells ml^-1^ in NH_4_ ^+^-YBCII using the correlation between OD_600_ and cell density. The suspensions were then serially diluted to obtain cell densities of 10^7^, 10^6^, and 10^5^ cells ml^-1^. Liquid cultures of *M. litorale, M. mangrovicola* and *M. rhizophilum* were adjusted to 10^7^ cells ml^-1^ in NH_4_ ^+^-YBCII using the correlation between OD_600_ and cell density. Liquid cultures of *V. diazotrophicus* were adjusted to 10^7^ cells ml^-1^ in YBCII using the correlation between OD_600_ and cell density. The suspensions were then serially diluted to obtain cell densities of 10^6^, 10^5^, and 10^4^ cells ml^-1^. The gDNA from the dilutions was then extracted as described above and quantified. The resulting calibration curves were used to correlate Ct values with cell densities (Extended Data Fig. 6b). gDNA was quantified by qPCR using the Luna Universal qPCR Master Mix (New England Biolabs) on an AriaMx Real-time PCR System (Agilent) according to the manufacturer’s protocol. No template control samples and no amplification control samples were included in each run. After baseline correction, the Ct values of the samples were obtained. The following organism-specific primers (5’ to 3’) were used that do not cross-amplify the other organisms: GTGTAGGGCAAAGAGGGACC and GGACTTGGTGGAGTGATGGG for *Chlamydomonas* sp., GAATCTGCCTGGTAGTGGGG and AGCTACGGATCATCGCCTTG for *M. stanieri*, AAGAAGCACCGGCTAACTCC and CGGGGCTTTCACATCTGACT for *M. litorale*, ATGCAAGTCGAGCGGTAACA and GAAGAGCCCCCACTTTCCTC for *M. mangrovicola*, GCTAATACCGCATACGCCCT and AGCTAAGGATCGTCGCCTTG for *M. rhizophilum*, GGAGAAAGCAGGGGACCTTC and GGACCGTGTCTCAGTTCCAG for *P. deceptionensis* and TCGTGAGGAAGGTGGTGTTG and CCGGGCTTTCACATCTGACT for *V. diazotrophicus*. The gDNAs standards extracts of 10^5^ cells ml^-1^ for *Chlamydomonas* sp. and 10^7^ cells ml^-1^ for the bacteria were used as internal controls in all qPCR rounds.

### Ammonium content

1 ml of cultures was collected at LD 6 on days 0, 4, 7, 11, 14, 18, and 21, and the cells were pelleted by centrifugation (5 min, 4000 x *g*, room temperature). Ammonium quantification was done with the supernatants using a commercially available colourimetric reaction kit (MQuant Ammonium [NH_4_ ^+^] test; Merck, Darmstadt, Germany).

### Chlorophyll and carotenoid quantifications

Pigment extraction and measurements were done at LD6 as described in Duanmu et al. (2013) and Minocha et al. (2009). Briefly, 1 ml of cell culture was collected on days 0, 4, 7, 11, 14, 18, and 21, and cells were pelleted by centrifugation. Pellets were dissolved in 1 ml of N, N-dimethylformamide (DMF), thoroughly vortexed, and then centrifuged again. The supernatants were measured at different wavelengths with the following formulae used for quantification:

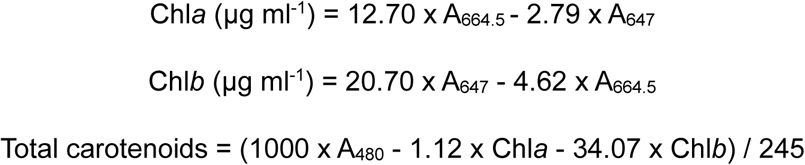

### Photosynthetic efficiency

For fluorescence measurements, cultures were grown in NH_4_ ^+^-YBCII media or bacterial spent media of *M. stanieri* (ΔSM_Msta) under an LD 12 h:12 h cycle as described above. On day 7 (LD 6) and before measurements, samples were diluted to a final concentration of 5 μg chlorophyll ml^-1^. A volume of 0.3 ml was placed into a KS-2500 suspension cuvette, where cells were dark-adapted for 30 min as described previously (Rredhi et al., 2021). The maximum quantum yield of PSII (Fv/Fm) was measured using a MINI-PAM-II (Heinz Walz GmbH, Germany) according to the manufacturer’s instructions.

### Electron microscopy (EM)

For ultrathin-section transmission EM, 10 ml of cells grown in fresh NH_4_^+^-YBCII media or ΔSM_Msta were harvested at LD6 on day 7 by centrifugation at 500 x *g*, and pre-fixed using 2.5% glutaraldehyde. Sample preparation followed the protocol described by Lübben et al. (2024). High-pressure freezing was carried out with an EM HPM100 system (Leica Microsystems, Wetzlar, Germany). This was followed by freeze-substitution in an EM AFS2 unit (Leica Microsystems) using a solution containing 0.2% osmium tetroxide, 0.1% uranyl acetate, and 9% water in pure acetone for 42 hours, as described by Peschke et al. (2013). Afterwards, samples were embedded in Epon 812 resin and polymerised at 63 °C. Ultrathin sections of approximately 70 nm thickness were cut using a 35° diamond knife (DiATOME, Nidau, Switzerland) on an Ultracut E ultramicrotome (Leica Microsystems). The sections were placed on collodion-coated 75-mesh copper grids (Science Services GmbH, Munich, Germany), post-stained with 80 mM lead citrate in NaOH at pH 13 and subsequently examined with an EM 912 transmission electron microscope (Zeiss, Oberkochen, Germany). The microscope was equipped with an integrated OMEGA energy filter operating in zero-loss mode at 80 kV. Images were recorded using a 2k × 2k slow-scan CCD camera (Tröndle Restlichtverstärkersysteme, Moorenweis, Germany).

### Starch assay

The starch assay was adapted from Vuong et al. (2025). 10 ml of algal culture (day 7, LD 6) was washed twice with PBS and then resuspended in 2 ml of PBS. Cells were then harvested by centrifugation for 5 min at 4000 x *g* at 23 °C. The supernatant was removed, and the cell pellet was frozen in liquid nitrogen. The cell pellet was resuspended in 1 ml of 80% (v/v) ethanol and incubated for 15 min at 80 °C and 1,200 rpm in a thermoshaker. The sample was centrifuged for 15 min at 16,000 x *g* at 4 °C, and the supernatant was discarded. 1 ml of ice-cold H_2_O was added to the pellet, and the samples were vortexed for 5 min. The supernatant was discarded, and the pellet was dried for 1 h in a speed-vac. 250 μL of 0.2 M KOH solution was added to the dried pellet, and the samples were heated to 95 °C under shaking at 1,200 rpm for 1 h. This step destroys the 3-D structure of the starch and brings it into solution. The reaction was neutralised by adding 42 μL of 1 M acetic acid, and the sample was centrifuged again. 200 μL of each sample was recovered for the starch assay.

The starch content of the cells was determined using the Enzytec™ Generic Starch kit (no. E1266, R-Biopharm AG, Darmstadt, Germany). 50 μL amyloglucosidase (AGS) solution was added to 200 μL of the sample. The reactions were incubated overnight at 55 °C. For the measurements, 200 μL of the overnight AGS reaction was mixed with 600 μL of kit-based reaction buffer in a microplate. The glucose content was determined after the addition of hexokinase/glucose-6-phosphate dehydrogenase by measuring the increase in absorbance at 340 nm, corresponding to the reduction of NADP+ to NADPH, H^+^ (Findinier et al., 2017) using a microplate reader (TECAN).

### Lipid assay and microscopy

Lipid droplet staining was performed according to Bensalem et al. (2018) with some modifications, by using BODIPY™ 505/515 (D3921, Invitrogen). BODIPY ™ was dissolved in DMSO to a final concentration of 0.1 mg ml ^-1^. At LD6 on day 7, 500 μL of the cell samples were taken, fixed in 4% (v/v) formalin for 10 min, and centrifuged for 5 min at 4000 x *g*. Cells were then incubated in BODIPY™ solutions at a final concentration of 1.5 μg ml^-1^ for 45 min in the dark. The samples were washed twice with NH_4_ ^+^-YBCII medium and resuspended in NH_4_ ^+^-YBCII before visualisation under the microscope.

For microscopy of lipid droplet staining, the Zeiss Elyra 7 was used in Apotome mode to suppress out-of-focus light. All images have been recorded using a Plan-Apochromat 40×/1.4 Oil DIC M27 objective. Illumination for the lipid stain imaging was provided by the 488 nm laser (set to 3-5%), and a 525/50 bandpass was used as an emission filter. For each imaged area, a z-stack was then recorded (approximately 8 μm total thickness) to image the complete cell. The raw apotome data were reconstructed with default settings using ZEN blue 3.11 for a final z-stepping of 0.11 μm. To facilitate the evaluation of the lipid stain, the stack was flattened using the maximum intensity projection function of ZEN blue.

### Statistical analyses

Data were analysed statistically and plotted in Microsoft Excel (Version 2604). Comparisons were conducted with Student’s *t*-test (2-tailed, unpaired). Data are representative of at least two independent experiments, and include routinely three biological replicates, unless otherwise indicated.

## Acknowledgements

We thank David Carrasco Flores, Vivien Hotter, Patrick Schwartze, Prateek Shetty and Teresa Zeibig for valuable discussions. We thank Prateek Shetty further for proofreading the manuscript. M.M. received funding from the Deutsche Forschungsgemeinschaft (DFG, German Research Foundation) within the collaborative research center SFB1127/2 ChemBioSys – Project ID 239748522, subproject A02. MM and PT received funding from the DFG within the Germany’s Excellence Strategy – EXC 2051 – Project ID 390713860. We thank the Microverse Imaging Center also financed by EXC 2051 for providing microscope facility support for data acquisition. MB was supported by ERC CoG EXPAND 101170891. The Free State of Thuringia funded the ELYRA 7 with grant no. 2019 FGI 0003.

## Author contributions

BBD, MO, MB and MM designed the project. BBD, MO, TV, PT and TY performed experiments, and all authors contributed to the analysis and interpretation of results. BBD and MM wrote the paper with input from all coauthors.

## Data availability

The data supporting this study’s findings have been included in this manuscript.

## Additional information

Extended data: Figures 1 to 6 and Tables 1 to 2

Extended Data Fig. 1 | Overview of *Chlamydomonas* sp. (Csp) mono- and cocultures with *V. diazotrophicus* (Vdiaz)

Extended Data Fig. 2 | Several *Marinobacterium* spp. interact mutualistically with *Chlamydomonas* sp. (Csp), in contrast to *P. deceptionensis* (Pdec)

Extended Data Fig. 3 | Small heat-resistant compounds produced by *M. stanieri* (Msta) in mono- and cocultures improve *Chlamydomonas* sp. (Csp) growth, while the bacterium benefits from small heat-resistant compounds produced during cocultures only

Extended Data Fig. 4 | Heat-resistant bacterial metabolite(s) increase photosynthetic activities in algal metabolism

Extended Data Fig. 5 | Benefits from bi- and tripartite interactions of *Chlamydomonas* sp. (Csp) with *M. stanieri* (Msta) and *V. diazotrophicus* (Vdiaz)

Extended Data Fig. 6 | Methods

Extended Data Table 1 | Statistical significances of data in Fig. 1 to Fig. 5

Extended Data Table 2 | Statistical significances of data in Extended Data Fig. 2 to Extended Data Fig. 5

## References

1. Aiyar, P., Schaeme, D., García-Altares, M., Carrasco Flores, D., Dathe, H., Hertweck, C., et al. (2017). Antagonistic bacteria disrupt calcium homeostasis and immobilize algal cells. Nat. Commun. 8, 1756. doi: 10.1038/s41467-017-01547-8

2. Allen, M. D., Del Campo, J. A., Kropat, J., and Merchant, S. S. (2007). *FEA1*, *FEA2*, and *FRE1*, encoding two homologous secreted proteins and a candidate ferrireductase, are expressed coordinately with *FOX1* and *FTR1* in iron-deficient *Chlamydomonas reinhardtii*. Eukaryot. Cell 6, 1841–1852. doi: 10.1128/EC.00205-07

3. Baumann, P., Bowditch, R. D., Baumann, L., and Beaman, B. (1983). Taxonomy of marine *Pseudomonas* species: *P. stanieri* sp. nov.; *P. perfectomarina* sp. nov., nom. rev.; *P. nautica*; and *P. doudoroffii*. Int. J. Syst. Bacteriol. 33, 857–865. doi: 10.1099/00207713-33-4-857

4. Bell, W., and Mitchell, R. (1972). Chemotactic and growth responses of marine bacteria to algal extracellular products. Biol. Bull. 143, 265–277. doi: 10.2307/1540052

5. Bensalem, S., Lopes, F., Bodénès, P., Pareau, D., Français, O., and Le Pioufle, B. (2018). Structural changes of *Chlamydomonas reinhardtii* cells during lipid enrichment and after solvent exposure. Data Brief 17, 1283–1287. doi: 10.1016/j.dib.2018.02.042

6. Bonnet, S., Benavides, M., Le Moigne, F. A. C., Camps, M., Torremocha, A., Grosso, O., et al. (2022). Diazotrophs are overlooked contributors to carbon and nitrogen export to the deep ocean. ISME J. 17, 47–58. doi: 10.1038/s41396-022-01319-3

7. Burgunter-Delamare, B., Shetty, P., Vuong, T., and Mittag, M. (2024). Exchange or eliminate: the secrets of algal-bacterial relationships. Plants 13, 829. doi: 10.3390/plants13060829

8. Calatrava, V., Hom, E. F. Y., Guan, Q., Llamas, A., Fernández, E., and Galván, A. (2024). Genetic evidence for algal auxin production in *Chlamydomonas* and its role in algal-bacterial mutualism. iScience 27, 108762. doi: 10.1016/j.isci.2023.108762

9. Carrasco Flores, D., Fricke, M., Wesp, V., Desirò, D., Kniewasser, A., Hölzer, M., et al. (2021). A marine *Chlamydomonas* sp. emerging as an algal model. J. Phycol. 57, 54–69. doi: 10.1111/jpy.13083

10. Carrasco Flores, D., Hotter, V., Vuong, T., Hou, Y., Bando, Y., Scherlach, K., et al. (2024). A mutualistic bacterium rescues a green alga from an antagonist. Proc. Natl. Acad. Sci. 121, e2401632121. doi: 10.1073/pnas.2401632121

11. Chen, Y.-B., Zehr, J. P., and Mellon, M. (1996). Growth and nitrogen fixation of the diazotrophic filamentous nonheterocystous cyanobacterium *Trichodesmium* sp. ims 101 in defined media: evidence for a circadian rhythm. J. Phycol. 32, 916–923. doi: 10.1111/j.0022-3646.1996.00916.x

12. Cirri, E., and Pohnert, G. (2019). Algae-bacteria interactions that balance the planktonic microbiome. New Phytol. 223, 100–106. doi: 10.1111/nph.15765

13. Cooper, M. B., Kazamia, E., Helliwell, K. E., Kudahl, U. J., Sayer, A., Wheeler, G. L., et al. (2019). Cross-exchange of B-vitamins underpins a mutualistic interaction between *Ostreococcus tauri* and *Dinoroseobacter shibae*. ISME J. 13, 334–345. doi: 10.1038/s41396-018-0274-y

14. Crétin, P., Mahoudeau, L., Joublin-Delavat, A., Paulhan, N., Labrune, E., Verdon, J., et al. (2025). High metabolic versatility and phenotypic heterogeneity in a marine non-cyanobacterial diazotroph. Curr. Biol. 35, 2659–2671.e3. doi: 10.1016/j.cub.2025.04.071

15. Croft, M. T., Lawrence, A. D., Raux-Deery, E., Warren, M. J., and Smith, A. G. (2005). Algae acquire vitamin B_12_ through a symbiotic relationship with bacteria. Nature 438, 90–93. doi: 10.1038/nature04056

16. De Hostos, E. L., Schilling, J., and Grossman, A. R. (1989). Structure and expression of the gene encoding the periplasmic arylsulfatase of *Chlamydomonas reinhardtii*. Mol. Gen. Genet. MGG 218, 229–239. doi: 10.1007/BF00331273

17. Delaux, P.-M., and Gutjahr, C. (2024). Evolution of small molecule-mediated regulation of arbuscular mycorrhiza symbiosis. Philos. Trans. R. Soc. B Biol. Sci. 379, 20230369. doi: 10.1098/rstb.2023.0369

18. Deng, Y., Mauri, M., Vallet, M., Staudinger, M., Allen, R. J., and Pohnert, G. (2022). Dynamic diatom-bacteria consortia in synthetic plankton communities. Appl. Environ. Microbiol. 88, e01619–22. doi: 10.1128/aem.01619-22

19. Dixon, R., and Kahn, D. (2004). Genetic regulation of biological nitrogen fixation. Nat. Rev. Microbiol. 2, 621–631. doi: 10.1038/nrmicro954

20. Duanmu, D., Casero, D., Dent, R. M., Gallaher, S., Yang, W., Rockwell, N. C., et al. (2013). Retrograde bilin signaling enables *Chlamydomonas* greening and phototrophic survival. Proc. Natl. Acad. Sci. 110, 3621–3626. doi: 10.1073/pnas.1222375110

21. Duncan, A., Barry, K., Daum, C., Eloe-Fadrosh, E., Roux, S., Schmidt, K., et al. (2022). Metagenome-assembled genomes of phytoplankton microbiomes from the Arctic and Atlantic Oceans. Microbiome 10, 67. doi: 10.1186/s40168-022-01254-7

22. Fei, C., Wilson, A. T., Mangan, N. M., Wingreen, N. S., and Jonikas, M. C. (2022). Modelling the pyrenoid-based CO_2_-concentrating mechanism provides insights into its operating principles and a roadmap for its engineering into crops. Nat. Plants 8, 583–595. doi: 10.1038/s41477-022-01153-7

23. Field, C. B., Behrenfeld, M. J., Randerson, J. T., and Falkowski, P. (1998). Primary production of the biosphere: integrating terrestrial and oceanic components. Science 281, 237– 240. doi: 10.1126/science.281.5374.237

24. Findinier, J., Tunçay, H., Schulz-Raffelt, M., Deschamps, P., Spriet, C., Lacroix, J.-M., et al. (2017). The *Chlamydomonas mex1* mutant shows impaired starch mobilization without maltose accumulation. J. Exp. Bot. 68, 5177–5189. doi: 10.1093/jxb/erx343

25. Focardi, A., Bramucci, A. R., Ajani, P., Khalil, A., Raina, J.-B., and Seymour, J. R. (2025). Defining the ecological strategies of phytoplankton associated bacteria. Nat. Commun. 16, 6363. doi: 10.1038/s41467-025-61523-5

26. Fu, H., Smith, C. B., Sharma, S., and Moran, M. A. (2020). Genome sequences and metagenome-assembled genome sequences of microbial communities enriched on phytoplankton exometabolites. Microbiol. Resour. Announc. 9, e00724–20. doi: 10.1128/MRA.00724-20

27. Fulweiler, R. W., Rinehart, S., Taylor, J., Kelly, M. C., Berberich, M. E., Ray, N. E., et al. (2025). Global importance of nitrogen fixation across inland and coastal waters. Science 388, 1205–1209. doi: 10.1126/science.adt1511

28. Harrison, E., Meeda, Y., Gaikwad, T., Wheeler, G., and Helliwell, K. (2025). Nitrogen status exerts dynamic control over phosphorus sensing and acquisition via PSR1 in colimited marine diatoms. Sci. Adv. 11, eadw8260. doi: 10.1126/sciadv.adw8260

29. Henshaw, R. J., Moon, J., Stehnach, M. R., Bowen, B. P., Kosina, S. M., Northen, T. R., et al. (2024). Metabolites from intact phage-infected *Synechococcus* chemotactically attract heterotrophic marine bacteria. Nat. Microbiol. 9, 3184–3195. doi: 10.1038/s41564-024-01843-2

30. Herrera-Ardila, Y. M., Orrego, D., Bejarano-López, A. F., and Klotz-Ceberio, B. (2022). Effect of heat treatment on vitamin content during the manufacture of food products at industrial scale. DYNA 89, 127–132. doi: 10.15446/dyna.v89n223.99775

31. Ji, X., Verspagen, J. M. H., Stomp, M., and Huisman, J. (2017). Competition between cyanobacteria and green algae at low *versus* elevated CO_2_: who will win, and why? J. Exp. Bot. 68, 3815–3828. doi: 10.1093/jxb/erx027

32. Kirk, D. L. (2005). A twelve-step program for evolving multicellularity and a division of labor. BioEssays 27, 299–310. doi: 10.1002/bies.20197

33. Lipsman, V., Shlakhter, O., Rocha, J., and Segev, E. (2024). Bacteria contribute exopolysaccharides to an algal-bacterial joint extracellular matrix. Npj Biofilms Microbiomes 10, 36. doi: 10.1038/s41522-024-00510-y

34. Lübben, M. K., Klingl, A., Nickelsen, J., and Ostermeier, M. (2024). CLEM, a universal tool for analyzing structural organization in thylakoid membranes. Physiol. Plant. 176, e14417. doi: 10.1111/ppl.14417

35. Mackinder, L. C. M., Chen, C., Leib, R. D., Patena, W., Blum, S. R., Rodman, M., et al. (2017). A spatial interactome reveals the protein organization of the algal CO_2_-concentrating mechanism. Cell 171, 133–147.e14. doi: 10.1016/j.cell.2017.08.044

36. Mahoudeau, L., Crétin, P., Joublin-Delavat, A., Rodrigues, S., Guillouche, C., Louvet, I., et al. (2026). The interplay between the marine diazotroph *Vibrio diazotrophicus* and its prophage shapes both biofilm structure and nitrogen release. Appl. Environ. Microbiol. 92, e01564–25. doi: 10.1128/aem.01564-25

37. Minocha, R., Martinez, G., Lyons, B., and Long, S. (2009). Development of a standardized methodology for quantifying total chlorophyll and carotenoids from foliage of hardwood and conifer tree species. Can. J. For. Res. 39, 849–861. doi: 10.1139/X09-015

38. Moore, C. M., Mills, M. M., Arrigo, K. R., Berman-Frank, I., Bopp, L., Boyd, P. W., et al. (2013). Processes and patterns of oceanic nutrient limitation. Nat. Geosci. 6, 701–710. doi: 10.1038/ngeo1765

39. Moran, M. A., Kujawinski, E. B., Schroer, W. F., Amin, S. A., Bates, N. R., Bertrand, E. M., et al. (2022). Microbial metabolites in the marine carbon cycle. Nat. Microbiol. 7, 508– 523. doi: 10.1038/s41564-022-01090-3

40. Peschke, M., Moog, D., Klingl, A., Maier, U. G., and Hempel, F. (2013). Evidence for glycoprotein transport into complex plastids. Proc. Natl. Acad. Sci. 110, 10860–10865. doi: 10.1073/pnas.1301945110

41. Rredhi, A., Petersen, J., Schubert, M., Li, W., Oldemeyer, S., Li, W., et al. (2021). DASH cryptochrome 1, a UV-A receptor, balances the photosynthetic machinery of *Chlamydomonas reinhardtii*. New Phytol. 232, 610–624. doi: 10.1111/nph.17603

42. Sanz-Luque, E., Chamizo-Ampudia, A., Llamas, A., Galvan, A., and Fernandez, E. (2015). Understanding nitrate assimilation and its regulation in microalgae. Front. Plant Sci. 6, 899. doi: 10.3389/fpls.2015.00899

43. Sasso, S., Stibor, H., Mittag, M., and Grossman, A. R. (2018). From molecular manipulation of domesticated *Chlamydomonas reinhardtii* to survival in nature. eLife 7, e39233. doi: 10.7554/eLife.39233

44. Satomi, M., Kimura, B., Hamada, T., Harayama, S., and Fujii, T. (2002). Phylogenetic study of the genus *Oceanospirillum* based on 16S rRNA and gyrB genes: emended description of the genus *Oceanospirillum*, description of *Pseudospirillum* gen. nov., *Oceanobacter* gen. nov. and *Terasakiella* gen. nov. and transfer of *Oceanospirillum jannaschii* and *Pseudomonas stanieri* to *Marinobacterium* as *Marinobacterium jannaschii* comb. nov. and *Marinobacterium stanieri* comb. no. Int. J. Syst. Evol. Microbiol. 52, 739–747. doi: 10.1099/00207713-52-3-739

45. Schmollinger, S., Mühlhaus, T., Boyle, N. R., Blaby, I. K., Casero, D., Mettler, T., et al. (2014). Nitrogen-sparing mechanisms in *Chlamydomonas* affect the transcriptome, the proteome, and photosynthetic metabolism. Plant Cell 26, 1410–1435. doi: 10.1105/tpc.113.122523

46. Schnellbaecher, A., Binder, D., Bellmaine, S., and Zimmer, A. (2019). Vitamins in cell culture media: stability and stabilization strategies. Biotechnol. Bioeng. 116, 1537–1555. doi: 10.1002/bit.26942

47. Seyedsayamdost, M. R., Case, R. J., Kolter, R., and Clardy, J. (2011). The Jekyll-and-Hyde chemistry of *Phaeobacter gallaeciensis*. Nat. Chem. 3, 331–335. doi: 10.1038/nchem.1002

48. Shimamura, D., Ikeuchi, T., Matsuda, A., Tsuji, Y., Fukuzawa, H., Mochida, K., et al. (2024). Periplasmic carbonic anhydrase CAH1 contributes to high inorganic carbon affinity in *Chlamydomonas reinhardtii*. Plant Physiol. 196, 2395–2404. doi: 10.1093/plphys/kiae463

49. Shortelle, A. B., and Colburn, E. A. (1987). Ecological effects of liming in a Cape Cod kettle pond: a note for fisheries managers. Lake Reserv. Manag. 3, 436–443. doi: 10.1080/07438148709354801

50. Vuong, T., Shetty, P., Kurtoglu, E., Schultz, C., Schrader, L., Then, P., et al. (2025). Metamorphosis of a unicellular green alga in the presence of acetate and a spatially structured three-dimensional environment. New Phytol. 245, 1180–1196. doi: 10.1111/nph.20299

51. Zehr, J. P., and Capone, D. G. (2020). Changing perspectives in marine nitrogen fixation. Science 368, eaay9514. doi: 10.1126/science.aay9514

52. Zehr, J. P., and Kudela, R. M. (2011). Nitrogen cycle of the open ocean: from genes to ecosystems. Annu. Rev. Mar. Sci. 3, 197–225. doi: 10.1146/annurev-marine-120709-142819

