## Supplemental Files for "Tripartite synergy - Metabolic crosstalk between two bacterial mutualists and a marine microalga promotes algal fitness"

### Extended Data

Extended Data Figure 1 to Figure 6: Page 2-10

Extended Data Table 1 to Table 2: Page 11-23

### Extended Data Figures

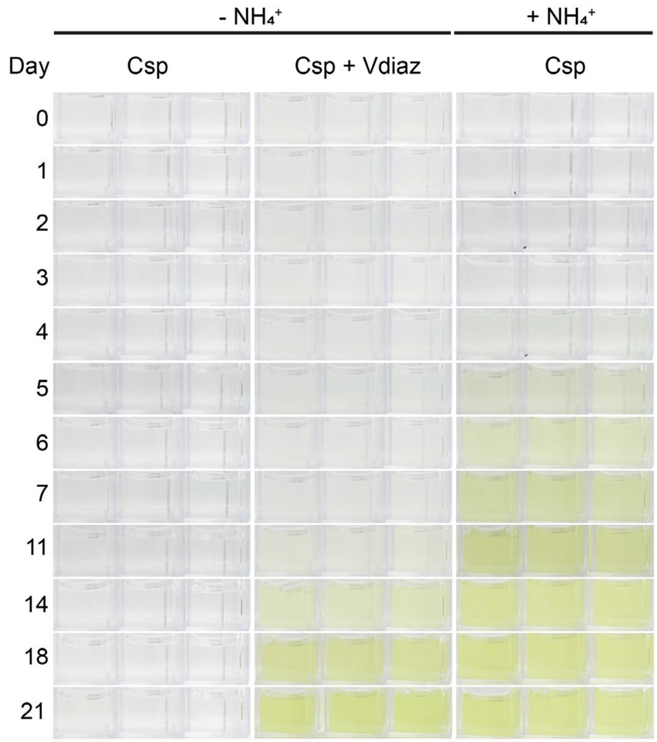


**Extended Data Fig. 1 | Overview of *Chlamydomonas* sp. (Csp) mono- and cocultures with *V. diazotrophicus* (Vdiaz).** The organisms were cocultivated in an $NH_4^+$-depleted medium (see Methods) over 21 days. An axenic culture of Csp in an $NH_4^+$-supplemented YBCII medium was used as a positive control. All experiments were done with $n = 3$ independent biological replicates.

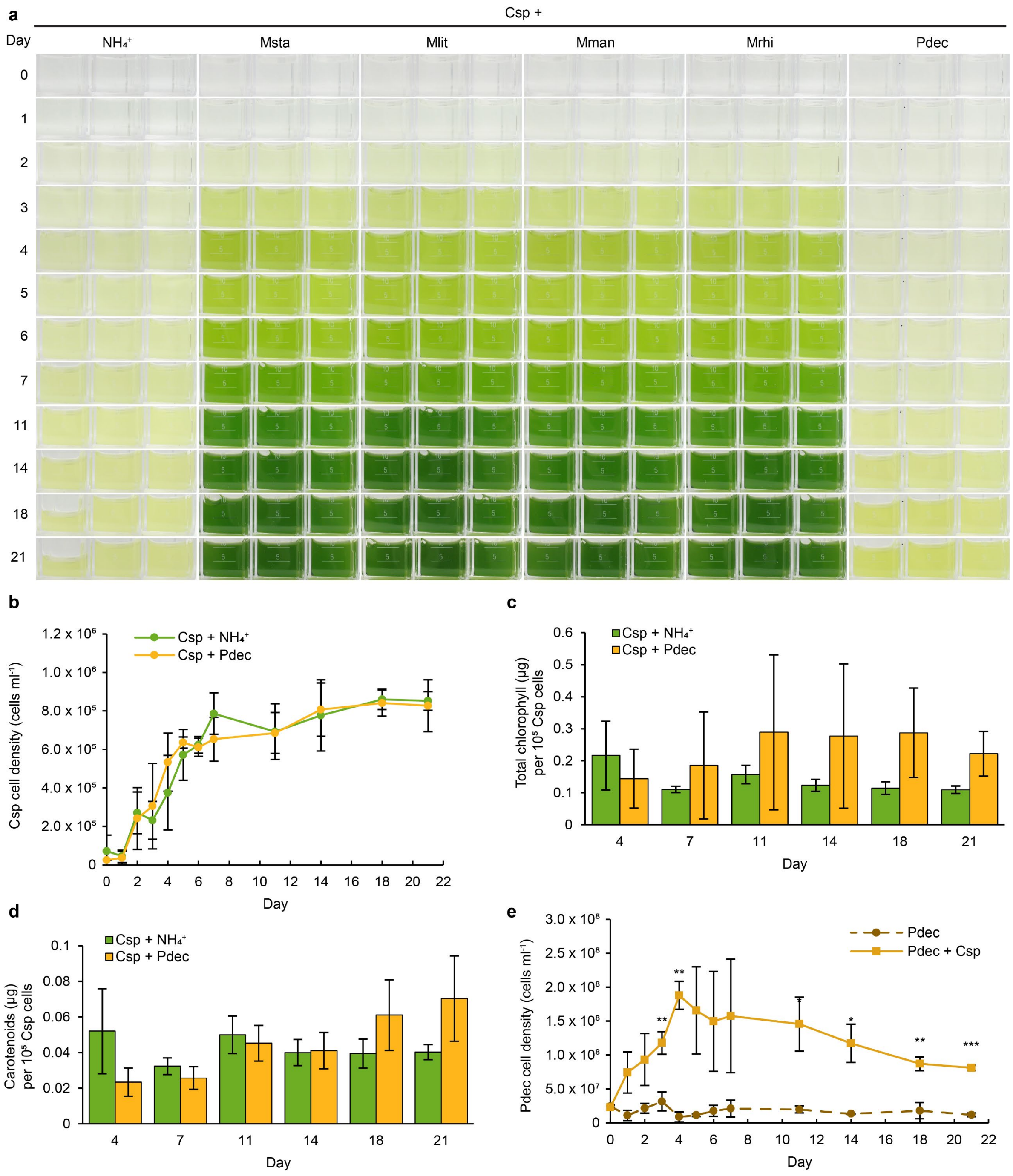


**Extended Data Fig. 2 | Several *Marinobacterium* spp. interact mutualistically with *Chlamydomonas* sp. (Csp), in contrast to *P. deceptionensis* (Pdec).** **a**, Overview of algal mono- and cocultures with *Marinobacterium* spp. Organisms were cultivated in an $NH_4^+$-supplemented YBCII medium over 21 days (see Methods). An axenic culture of Csp in an $NH_4^+$-supplemented medium was used as a positive control. Msta: *M. stanieri*, Mlit: *M. litorale*, Mman: *M. mangrovicola*, Mrhi: *M. rhizophilum*. **b**, Algal cell densities in mono- and cocultures with Pdec. For cultivation, see legend of (**a**). Cells from a 10 ml cell suspension were harvested over 21 days. Genomic DNA was extracted and used for qPCR to determine cell densities (Methods). **c-d**, Normalised total chlorophyll (**c**) and carotenoids (**d**) contents per $10^5$ cells in Csp mono- and cocultures with Pdec. **e**, Bacterial cell densities of Pdec in mono- and cocultures. For cultivation, see legend of (**a**). Cells from a 10 ml cell suspension were harvested over 21 days. Genomic DNA was extracted and used for qPCR to determine cell densities (Methods). **a-e**, All experiments were done with *n* = 3 independent biological replicates. Error bars indicate SDs. Asterisks indicate significant differences as calculated by Student's *t*-test: ***$P$ <0.001, **$P$ <0.01, *$P$ <0.05. Full statistical analyses are detailed in Extended Data Table 2a-d.

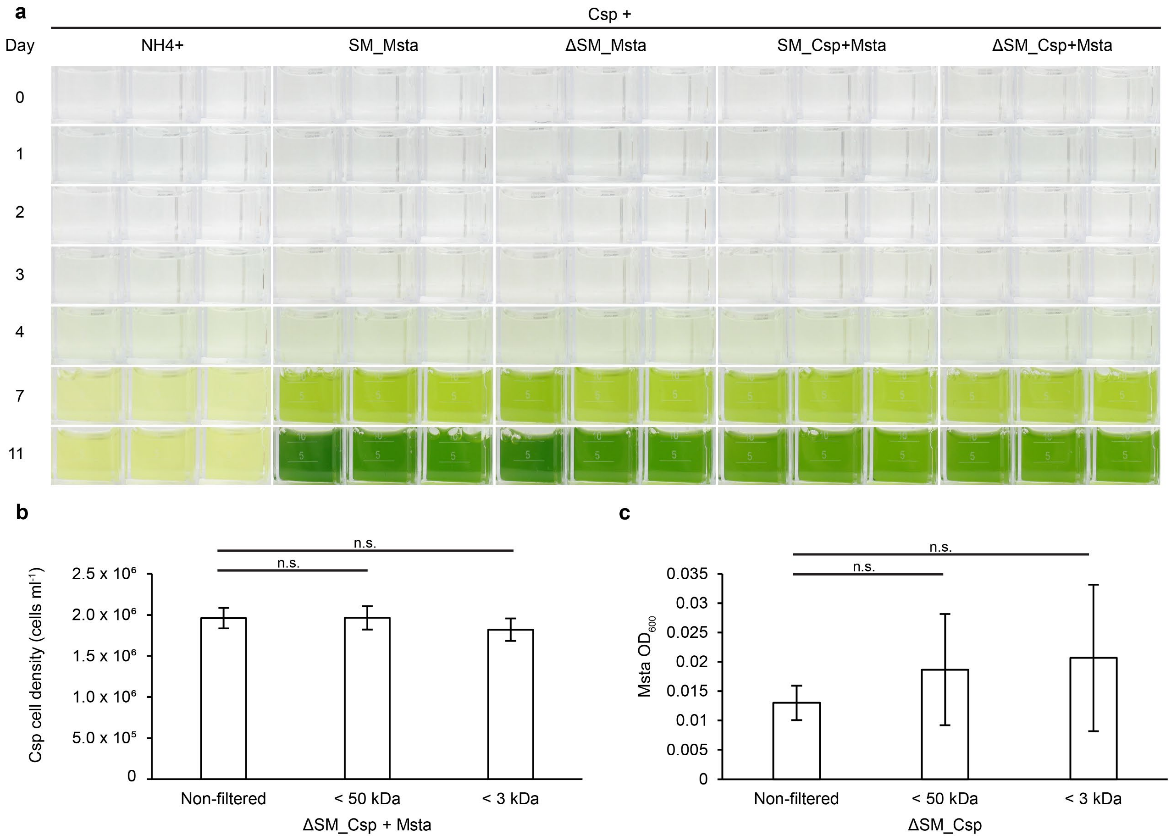


**Extended Data Fig. 3 | Small heat-resistant compounds produced by *M. stanieri* (Msta) in mono- and cocultures improve *Chlamydomonas* sp. (Csp) growth, while the bacterium benefits from small heat-resistant compounds produced during cocultures only.** **a**, Overview of algal monocultures in different spent media (SM). Csp was grown in spent media of Msta (SM_Msta) or in coculture spent media (SM_Csp+Msta) over 11 days. Heat-treated spent media (Methods) are indicated by "Δ" (**a-c**). Axenic cultures of Csp in $NH_4^+$-YBCII medium were used as a positive control. **b**, Algal cell densities after one week in non-filtered heated coculture spent media or after size fractionation (ΔSM_Csp+Msta). **b-c**, The shown statistics are compared with Csp grown in non-filtered ΔSM_Csp+Msta. **c**, Bacterial cell densities after one week in different fractions of ΔSM_Csp (see **b**). **a-c**, All experiments were done with $n$ = 3 independent biological replicates. **b-c,** Error bars indicate SDs. Asterisks indicate significant differences as calculated by Student's $t$-test: n.s., not significant. Full statistical analyses are detailed in Extended Data Table 2e-f.

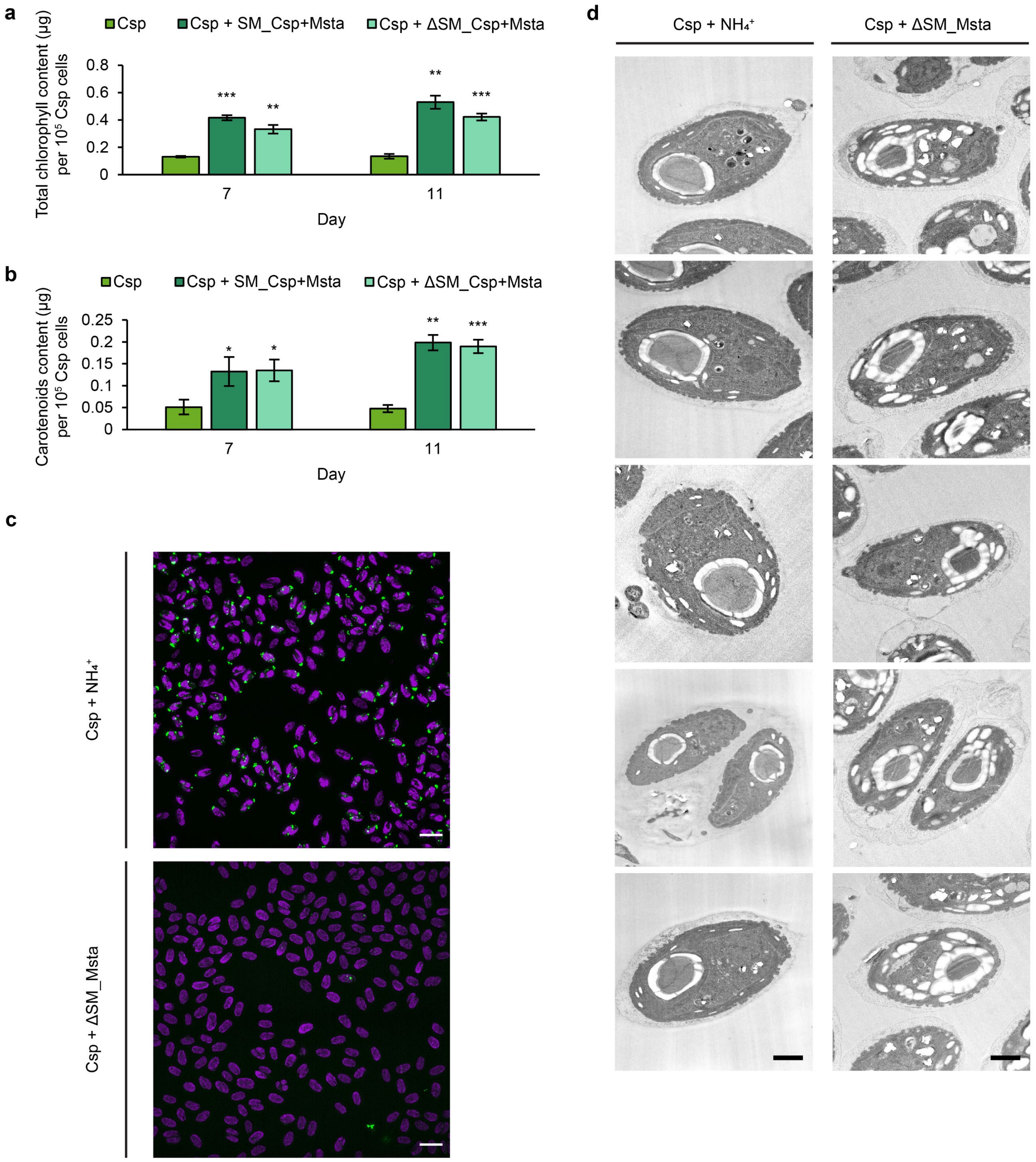


**Extended Data Fig. 4 | Heat-resistant bacterial metabolite(s) increase photosynthetic activities in algal metabolism. a-b**, Normalised total chlorophyll (**a**) and carotenoids (**b**) content per $10^5$ cells in *Chlamydomonas* sp. (Csp) cells grown in $NH_4^+$-YBCII, or in spent media of a Csp coculture with *M. stanieri* (SM_Csp+Msta) or in heated spent media (ΔSM_Csp+Msta). The statistics shown are in comparison with Csp grown in $NH_4^+$-YBCII. **c**, Electron micrographs of high-pressure frozen ultra-thin sections of Csp+$NH_4^+$ (left panels) and Csp+ΔSM_Msta (right panels). Scale bars = 1 µm. **d**, Stained lipid droplets (in green colour) and combined purple colored *Chl* autofluorescence of Csp+$NH_4^+$ (upper panel) and Csp+ΔSM_Msta (lower panel). Scale bars = 10 µm. **a-d**, Experiments were done with $n = 3$ (a-b-d) and $n = 2$ (c) independent biological replicates. **a-b**, Error bars indicate SDs. Asterisks indicate significant differences as calculated by Student's *t*-test: $^{***}P<0.001$, $^{**}P<0.01$, $^{*}P<0.05$. Full statistical analyses are detailed in Extended Data Table 2g-h.

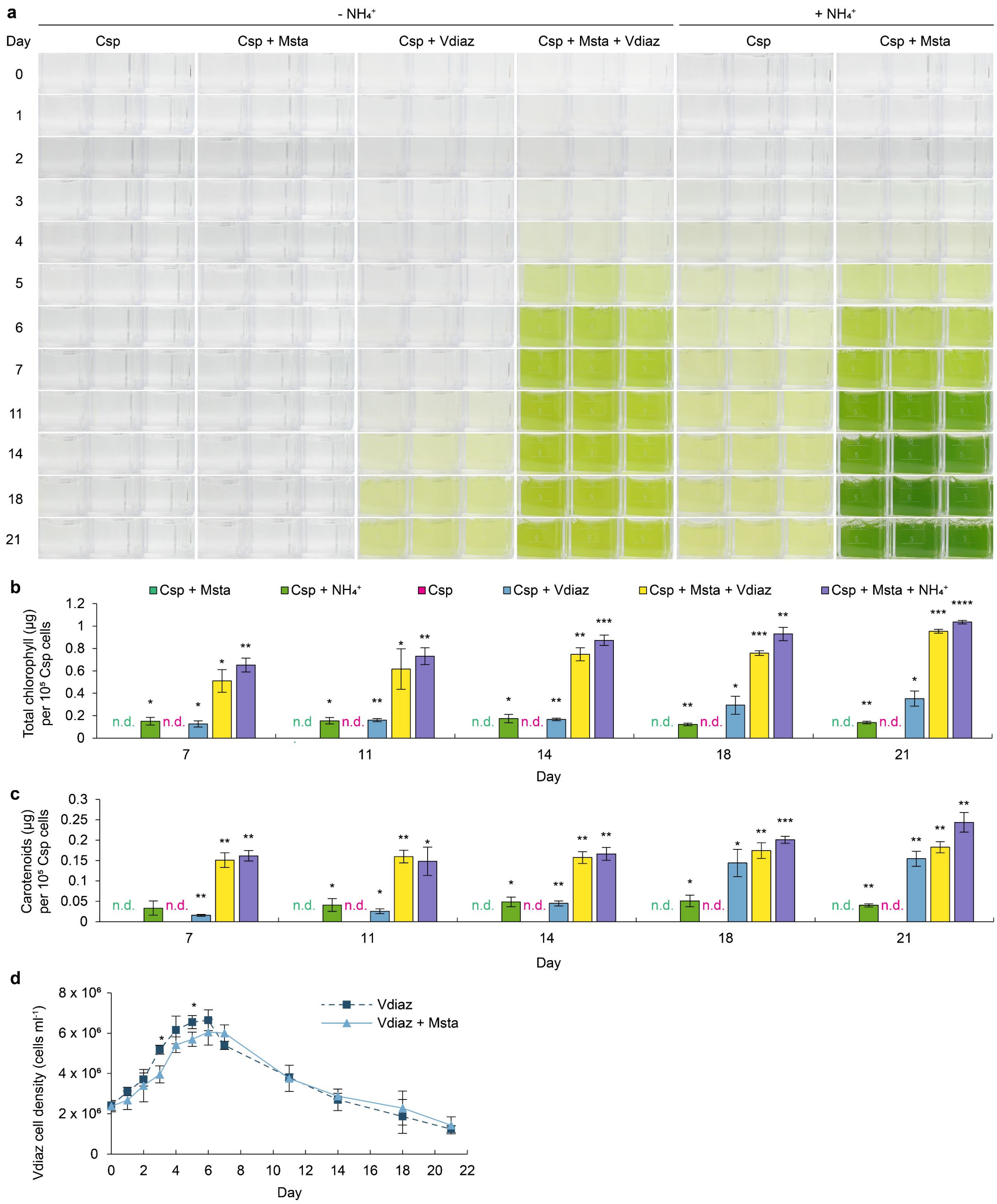


**Extended Data Fig. 5 | Benefits from bi- and tripartite interactions of *Chlamydomonas* sp. (Csp) with *M. stanieri* (Msta) and *V. diazotrophicus* (Vdiaz). a**, Overview of the tripartite cultures of Csp with Vdiaz and Msta. The organisms were cultivated in an $NH_4^+$-depleted YBCII medium over 21 days (see Methods). Csp+Msta cocultures in an $NH_4^+$-supplemented YBCII medium were used as a positive control. **b-c**, Normalised total chlorophyll (**b**) and carotenoids (**c**) content per $10^5$ cells in Csp cocultures. n.d.: not detected. The shown statistics are in comparison with Csp+Msta cocultures in $NH_4^+$-depleted YBCII medium. **d**, Bacterial cell densities of *Vdiaz* in mono- and cocultures with *Msta*. For cultivation, see legend of (**a**). Cells from a 10 ml cell suspension were harvested over 21 days. Genomic DNA was extracted and used for qPCR to determine cell densities (Methods). **a-d**, All experiments were done with

$n = 3$ independent biological replicates. **b-d**, Error bars indicate SDs. Asterisks indicate significant differences as calculated by Student's $t$-test: ****$P < 0.0001$, ***$P < 0.001$, **$P < 0.01$, *$P < 0.05$. Full statistical analyses are detailed in Extended Data Table 2i-k.

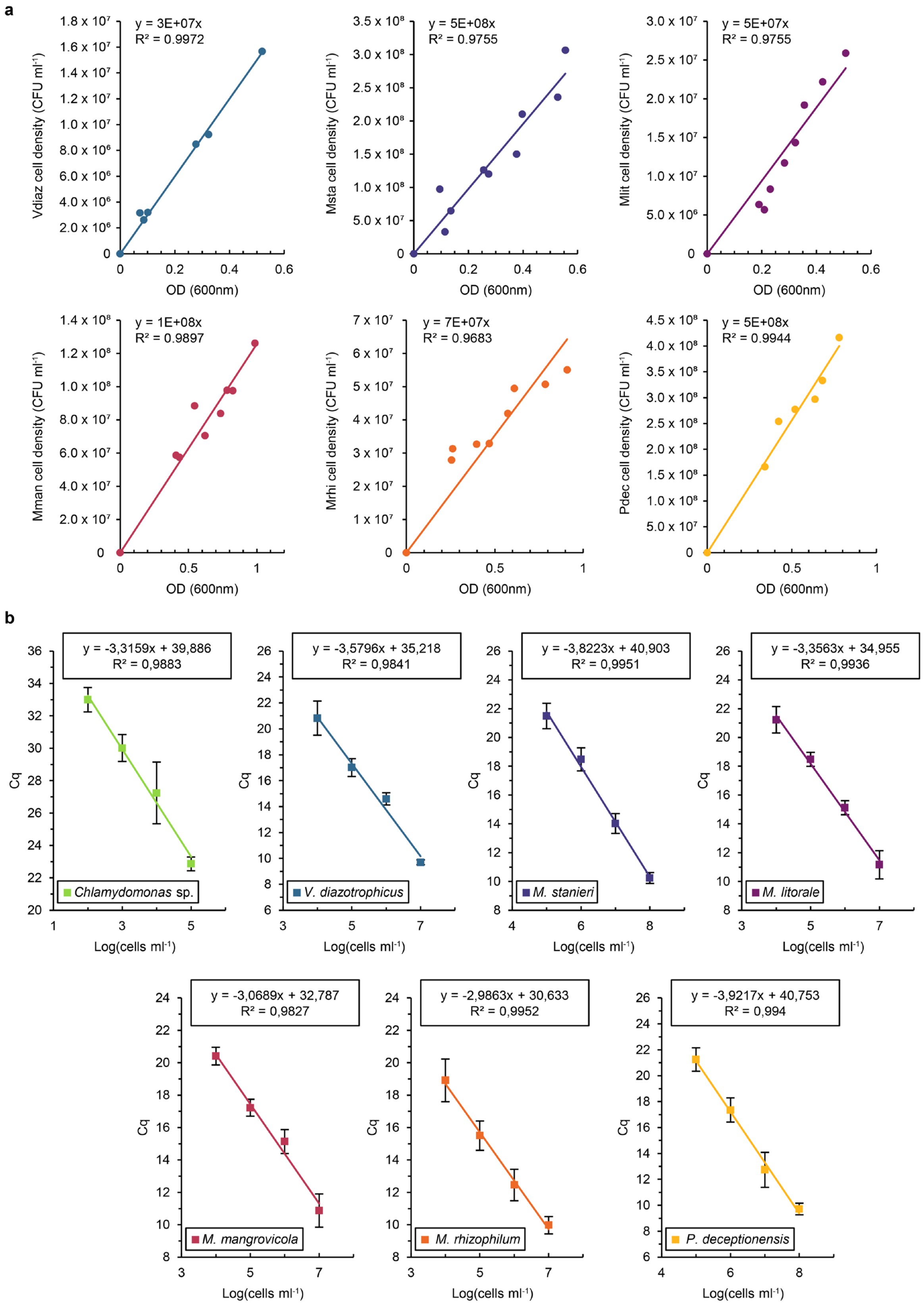


**Extended Data Fig. 6 | Methods. a**, Correlation of $OD_{600}$ values to bacterial cell densities (Vdiaz: *V. diazotrophicus*, Msta: *M. stanieri*, Mlit: *M. litorale*, Mman: *M. mangrovicola*, Mrhi: *M. rhizophilum*. Pdec: *P. deceptionensis*). **b**, Calibration curves for algal and bacterial cell densities based on genomic DNA quantification. Cell densities calculated from cell counts for

*Chlamydomonas* sp. or from $OD_{600}$ values for bacteria were correlated with the cycle threshold (Ct) of the qPCR. **a-b**, All experiments were done with $n = 3$ independent biological replicates. Error bars indicate SDs.

### Extended Data Tables

**Extended Data Table 1 | Statistical significances of data in Fig. 1 to Fig. 5.** Asterisks indicate significant differences as calculated by Student's *t*-test: ****$P$ <0.0001, ***$P$ <0.001, **$P$ <0.01, *$P$ <0.05. n.s.: not significant.

a) Fig.1c: p-values of cell densities for *Chlamydomonas* sp. (Csp) in mono and cocultures with *V. diazotrophicus* (Vdiaz) in an $NH_4^+$-depleted YBCII medium compared to Csp monocultures in an $NH_4^+$-supplemented YBCII medium.

| Day | Csp | Csp+Vdiaz |
|---|---|---|
| 0 | n.s. | n.s. |
| 1 | n.s. | n.s. |
| 2 | n.s. | n.s. |
| 3 | * | n.s. |
| 4 | * | * |
| 5 | *** | * |
| 6 | * | * |
| 7 | * | * |
| 11 | * | * |
| 14 | * | n.s. |
| 18 | ** | n.s. |
| 21 | ** | n.s. |

b) Fig. 1d: p-values of normalised $NH_4^+$ content in monocultures of *Chlamydomonas* sp. (Csp) and *V. diazotrophicus* (Vdiaz), compared to Csp+Vdiaz cocultures in an $NH_4^+$-depleted YBCII medium.

| Day | Csp | Vdiaz |
|---|---|---|
| 0 | n.s | n.s. |
| 4 | n.s. | n.s. |
| 7 | ** | n.s. |
| 11 | * | * |
| 14 | * | n.s. |
| 18 | n.s. | * |
| 21 | * | * |

Fig. 1d Inlet: p-values of total $NH_4^+$ content in monocultures of *Chlamydomonas* sp. (Csp) and *V. diazotrophicus* (Vdiaz), compared to Csp+Vdiaz cocultures in an $NH_4^+$-depleted YBCII medium.

| Day | Csp | Vdiaz |
|---|---|---|
| 0 | n.s | n.s. |
| 4 | n.s. | n.s. |
| 7 | ** | n.s. |
| 11 | * | n.s. |
| 14 | * | n.s. |
| 18 | n.s. | n.s. |
| 21 | * | * |

c) Fig. 1e: p-values of cell densities for *V. diazotrophicus* (Vdiaz) in cocultures with *Chlamydomonas* sp. (Csp), compared to Vdiaz monocultures in an $NH_4^+$-depleted YBCII medium.

| Day | Csp+Vdiaz |
|---|---|
| 0 | n.s. |
| 1 | n.s. |
| 2 | n.s. |
| 3 | n.s. |
| 4 | n.s |
| 5 | n.s |
| 6 | n.s |
| 7 | ** |
| 11 | ** |
| 14 | n.s. |
| 18 | * |
| 21 | n.s. |

d) Fig. 2b: p-values of cell densities for *Chlamydomonas* sp. (Csp) in cocultures with *Marinobacterium* spp. compared to monocultures of Csp. in an $NH_4^+$-supplemented YBCII medium. Msta: *M. stanieri*, Mlit: *M. litorale*, Mman: *M. mangrovicola*, and Mrhi: *M. rhizophilum.*

| Day | Csp+Msta | Csp+Mlit | Csp+Mman | Csp+Mrhi |
|---|---|---|---|---|
| 0 | n.s. | n.s. | n.s. | n.s. |

| | | | | |
|---|---|---|---|---|
| 1 | n.s. | n.s. | n.s. | n.s. |
| 2 | n.s. | n.s. | n.s. | n.s. |
| 3 | n.s. | * | n.s. | * |
| 4 | n.s. | n.s. | * | * |
| 5 | n.s. | * | n.s. | n.s. |
| 6 | n.s. | ** | n.s. | n.s. |
| 7 | ** | * | * | * |
| 11 | * | ** | * | * |
| 14 | **** | *** | ** | ** |
| 18 | ** | ** | ** | ** |
| 21 | * | * | ** | ** |

e) Fig. 2c: p-values of normalised total chlorophyll content of *Chlamydomonas* sp. (Csp) in cocultures with *Marinobacterium* spp. compared to Csp monoculture in an $NH_4^+$-supplemented YBCII medium. Msta: *M. stanieri*, Mlit: *M. litorale*, Mman: *M. mangrovicola*, and Mrhi: *M. rhizophilum*.

| Day | Csp+Msta | Csp+Mlit | Csp+Mman | Csp+Mrhi |
|---|---|---|---|---|
| 4 | n.s. | n.s. | n.s. | n.s. |
| 7 | ** | ** | * | * |
| 11 | ** | ** | ** | * |
| 14 | ** | ** | ** | ** |
| 18 | **** | ** | * | ** |
| 21 | ** | ** | ** | *** |

f) Fig. 2d: p-values of normalised carotenoid content of *Chlamydomonas* sp. (Csp) in cocultures with *Marinobacterium* spp., compared to Csp monocultures in an $NH_4^+$-supplemented YBCII medium. Msta: *M. stanieri*, Mlit: *M. litorale*, Mman: *M. mangrovicola*, Mrhi: *M. rhizophilum*.

| Day | Csp+Msta | Csp+Mlit | Csp+Mman | Csp+Mrhi |
|---|---|---|---|---|
| 4 | n.s. | n.s. | n.s. | n.s. |
| 7 | ** | * | * | * |
| 11 | n.s. | ** | ** | * |
| 14 | ** | *** | ** | * |
| 18 | *** | ** | ** | **** |
| 21 | ** | ** | **** | *** |

g) Fig. 2e: p-values of cell densities for *M. stanieri* (Msta) in cocultures with *Chlamydomonas* sp. (Csp) compared to Msta monocultures in an $NH_4^+$-supplemented YBCII medium.

| Day | Msta+Csp |
|---|---|
| 0 | n.s. |
| 1 | n.s. |
| 2 | n.s. |
| 3 | n.s. |
| 4 | * |
| 5 | * |
| 6 | ** |
| 7 | ** |
| 11 | * |
| 14 | * |
| 18 | * |
| 21 | ** |

h) Fig. 2f: p-values of cell densities for *M. rhizophilum* (Mrhi) in cocultures with *Chlamydomonas* sp. (Csp) compared to Mrhi monocultures in an $NH_4^+$-supplemented YBCII medium.

| Day | Mrhi+Csp |
|---|---|
| 0 | n.s. |
| 1 | n.s. |
| 2 | n.s. |
| 3 | n.s. |
| 4 | * |
| 5 | ** |
| 6 | ** |
| 7 | * |
| 11 | * |
| 14 | * |
| 18 | ** |
| 21 | * |

i) Fig. 2g: p-values of cell densities for *M. litorale* (Mlit) in cocultures with *Chlamydomonas* sp. (Csp) compared to Mlit monocultures in an $NH_4^+$-supplemented YBCII medium.

| Day | Mlit+Csp |
|---|---|
| 0 | n.s. |

| | |
|---|---|
| 1 | n.s. |
| 2 | n.s. |
| 3 | n.s. |
| 4 | n.s. |
| 5 | * |
| 6 | * |
| 7 | ** |
| 11 | ** |
| 14 | ** |
| 18 | ** |
| 21 | ** |

j) Fig. 2h: p-values of cell densities for *M. mangrovicola* (Mman) in cocultures with *Chlamydomonas* sp. (Csp) compared to Mman monocultures in an $NH_4^+$-supplemented YBCII medium.

| Day | Mman+Csp |
|---|---|
| 0 | n.s. |
| 1 | n.s. |
| 2 | * |
| 3 | n.s. |
| 4 | * |
| 5 | * |
| 6 | * |
| 7 | ** |
| 11 | *** |
| 14 | ** |
| 18 | * |
| 21 | ** |

k) Fig. 3b, c: p-values of cell densities for *Chlamydomonas* sp. (Csp) grown in different spent media compared to *Chlamydomonas* sp. grown in an $NH_4^+$-supplemented YBCII medium. SM_Msta: Bacterial spent media, ΔSM_Msta: Heat-treated bacterial spent media, SM_Csp+Msta: Cocultures spent media and ΔSM_Csp+Msta: Heat-treated cocultures spent media.

| Day | Csp+SM_Msta | Csp+ΔSM_Msta | Csp+SM_Csp+Msta | Csp+ΔSM_Csp+Msta |
|---|---|---|---|---|
| 0 | n.s. | n.s. | n.s. | n.s. |
| 1 | n.s. | n.s. | n.s. | n.s. |

| | | | | |
|---|---|---|---|---|
| 2 | n.s. | n.s. | * | n.s. |
| 3 | n.s. | n.s. | n.s. | n.s. |
| 4 | n.s. | n.s. | n.s. | n.s. |
| 7 | ** | *** | ** | *** |
| 11 | *** | *** | ** | *** |

l) Fig. 3d, e: p-values of cell densities for *M. stanieri* (Msta) grown in different spent media compared to *M. stanieri* grown in an $NH_4^+$-supplemented YBCII medium. SM_Csp: Algal spent media, ΔSM_Csp: Heat-treated algal spent media, SM_Csp+Msta: Cocultures spent media and ΔSM_Csp+Msta: Heat-treated cocultures spent media.

| Day | Msta+SM_Csp | Msta+ΔSM_Csp | Msta+SM_Csp+Msta | Msta+ΔSM_Csp+Msta |
|---|---|---|---|---|
| 0 | n.s. | n.s. | n.s. | n.s. |
| 1 | n.s. | n.s. | n.s. | n.s. |
| 2 | n.s. | n.s. | n.s. | * |
| 3 | n.s. | n.s. | n.s. | n.s. |
| 4 | n.s. | n.s. | *** | * |
| 7 | n.s. | n.s. | n.s. | * |
| 11 | n.s. | n.s. | * | ** |

m) Fig. 3f: p-values of cell densities for *Chlamydomonas* sp. (Csp) after one week in different fractions of ΔSM_Msta compared to Csp grown in non-filtered ΔSM_Msta.

| Day | < 50 kDa | < 3 kDa |
|---|---|---|
| 7 | n.s. | n.s. |

n) Fig. 3g: p-values of cell densities for *M. stanieri* (Msta) after one week in different fractions of ΔSM_Csp+Msta compared to Msta grown in non-filtered ΔSM_Csp+Msta.

| Day | < 50 kDa | < 3 kDa |
|---|---|---|
| 7 | n.s. | n.s. |

o) Fig. 4a: p-values of normalised total chlorophyll content for *Chlamydomonas* sp. (Csp) grown in different spent media compared to *Chlamydomonas* sp. grown in an $NH_4^+$-supplemented YBCII medium. SM_Msta: Bacterial spent media, ΔSM_Msta: Heat-treated bacterial spent media.

| Day | Csp+SM_Msta | Csp+ΔSM_Msta |
|---|---|---|

| | | |
|---|---|---|
| 7 | **** | *** |
| 11 | ** | *** |

p) Fig. 4b: p-values of normalised carotenoid content for *Chlamydomonas* sp. (Csp) grown in different spent media compared to *Chlamydomonas* sp. grown in an $NH_4^+$-supplemented YBCII medium. SM_Msta: Bacterial spent media, ΔSM_Msta: Heat-treated bacterial spent media.

| Day | Csp+SM_Msta | Csp+ΔSM_Msta |
|---|---|---|
| 7 | *** | ** |
| 11 | *** | ** |

q) Fig. 5b: p-values of cell densities for *Chlamydomonas* sp. (Csp) cells grown in coculture with *M. stanieri* (Msta) in an $NH_4^+$-supplemented YBCII medium or in co- and tripartite cultures in an $NH_4^+$-depleted YBCII medium, compared to Csp+Msta cocultures grown in $NH_4^+$-depleted YBCII medium. Vdiaz: *V. diazotrophicus*.

| Day | Csp+Msta+$NH_4^+$ | Csp+Vdiaz | Csp+Msta+Vdiaz |
|---|---|---|---|
| 0 | n.s. | n.s. | n.s. |
| 1 | n.s. | n.s. | n.s. |
| 2 | n.s. | n.s. | n.s. |
| 3 | n.s. | n.s. | n.s. |
| 4 | n.s. | n.s. | * |
| 5 | * | n.s. | * |
| 6 | * | * | * |
| 7 | ** | * | * |
| 11 | *** | ** | ** |
| 14 | ** | ** | * |
| 18 | * | * | ** |
| 21 | * | * | ** |

r) Fig. 5c: p-values of normalised total chlorophyll content of *Chlamydomonas* sp. (Csp) cells grown in coculture with *M. stanieri* (Msta) in an $NH_4^+$-supplemented YBCII medium or in co- and tripartite cultures in an $NH_4^+$-depleted YBCII medium, compared to Csp+Msta cocultures grown in $NH_4^+$-depleted YBCII medium. Vdiaz: *V. diazotrophicus*. n.d.: not detected.

| Day | Csp+Msta+$NH_4^+$ | Csp+Vdiaz | Csp+Msta+Vdiaz |
|---|---|---|---|

| | | | |
|---|---|---|---|
| 7 | ** | * | * |
| 21 | **** | * | *** |

s) Fig.5 d: p-values of normalised carotenoid content of *Chlamydomonas* sp. (Csp) cells grown in coculture with *M. stanieri* (Msta) in an $NH_4^+$-supplemented YBCII medium or in co- and tripartite cultures in an $NH_4^+$-depleted YBCII medium, compared to Csp+Msta cocultures grown in $NH_4^+$-depleted YBCII medium. Vdiaz: *V. diazotrophicus*. n.d.: not detected.

| Day | Csp+Msta+$NH_4^+$ | Csp+Vdiaz | Csp+Msta+Vdiaz |
|---|---|---|---|
| 7 | ** | ** | ** |
| 21 | ** | ** | ** |

t) Fig. 5e: p-values of cell densities of *V. diazotrophicus* (Vdiaz) in tripartite interaction compared to Vdiaz cocultures with *Chlamydomonas* sp. (Csp) in an $NH_4^+$-depleted YBCII medium. Msta: *M. stanieri*.

| Day | Vdiaz+Csp+Msta |
|---|---|
| 0 | n.s. |
| 1 | n.s. |
| 2 | n.s. |
| 3 | n.s. |
| 4 | n.s. |
| 5 | n.s. |
| 6 | *** |
| 7 | n.s. |
| 11 | n.s. |
| 14 | n.s. |
| 18 | n.s. |
| 21 | n.s. |

u) Fig. 5f: p-values of cell densities for *M. stanieri* (Msta) in cocultures with *V. diazotrophicus* (Vdiaz) compared to Msta in monocultures in an $NH_4^+$-depleted YBCII medium.

| Day | Msta+Vdiaz |
|---|---|
| 0 | n.s. |
| 1 | n.s. |
| 2 | n.s. |
| 3 | n.s. |

| | |
|---|---|
| 4 | n.s. |
| 5 | n.s. |
| 6 | * |
| 7 | ** |
| 11 | * |
| 14 | n.s. |
| 18 | ** |
| 21 | * |

v) Fig. 5g: p-values of normalised $NH_4^+$ content in mono-, co- and tripartite cultures of *Chlamydomonas* sp. (Csp), *M. stanieri* (Msta) and *V. diazotrophicus* (Vdiaz) compared to Vdiaz monocultures in an $NH_4^+$-depleted YBCII medium.

| Day | Csp | Msta | Csp+Msta | Msta+Vdiaz | Csp+Vdiaz | Csp+Msta+Vdiaz |
|---|---|---|---|---|---|---|
| 0 | n.s. | n.s. | n.s. | n.s. | n.s. | n.s. |
| 4 | ** | ** | ** | n.s. | n.s. | n.s. |
| 7 | *** | *** | *** | n.s. | n.s. | n.s. |
| 11 | *** | *** | *** | n.s. | * | n.s. |
| 14 | * | * | * | n.s. | * | n.s. |
| 18 | ** | ** | ** | n.s. | ** | ** |
| 21 | * | * | * | n.s. | * | * |

w) Fig. 5h: p-values of cell densities for *M. stanieri* (Msta) in cocultures with *Chlamydomonas* sp. (Csp) with or without $NH_4^+$, compared to tripartite cultures in an $NH_4^+$-depleted YBCII medium.

| Day | Msta+Csp | Msta+Csp+$NH_4^+$ |
|---|---|---|
| 0 | n.s. | n.s. |
| 1 | n.s. | n.s. |
| 2 | n.s. | n.s. |
| 3 | n.s. | n.s. |
| 4 | ** | n.s. |
| 5 | **** | n.s. |
| 6 | ** | n.s. |
| 7 | **** | n.s. |
| 11 | * | n.s. |
| 14 | **** | * |
| 18 | *** | n.s. |
| 21 | **** | n.s. |

**Extended Data Table 2 | Statistical significances of data in Extended Data Fig. 2 to Extended Data Fig. 5.** Asterisks indicate significant differences as calculated by Student's *t*-test: ****$P$<0.0001, ***$P$<0.001, **$P$<0.01, *$P$<0.05. n.s.: not significant.

a) Extended Data Fig. 2b: p-values of cell densities for *Chlamydomonas* sp. (Csp) in cocultures with *P. deceptionensis* (Pdec) compared to Csp monocultures in an $NH_4^+$-supplemented YBCII medium.

| Day | Csp+Pdec |
|---|---|
| 0 | n.s. |
| 1 | n.s. |
| 2 | n.s. |
| 3 | n.s. |
| 4 | n.s. |
| 5 | n.s. |
| 6 | n.s. |
| 7 | n.s. |
| 11 | n.s. |
| 14 | n.s. |
| 18 | n.s. |
| 21 | n.s. |

b) Extended Data Fig. 2c: p-values of normalised total chlorophyll content of *Chlamydomonas* sp. (Csp) in cocultures with *P. deceptionensis* (Pdec) compared to Csp monocultures in an $NH_4^+$-supplemented YBCII medium.

| Day | Csp+Pdec |
|---|---|
| 4 | n.s. |
| 7 | n.s. |
| 11 | n.s. |
| 14 | n.s. |
| 18 | n.s. |
| 21 | n.s. |

c) Extended Data Fig. 2d: p-values of normalised carotenoid content of *Chlamydomonas* sp. (Csp) in cocultures with *P. deceptionensis* (Pdec) compared to Csp monocultures in an $NH_4^+$-supplemented YBCII medium.

| Day | Csp+Pdec |
|---|---|
| 4 | n.s. |
| 7 | n.s. |

| | |
|---|---|
| 11 | n.s. |
| 14 | n.s. |
| 18 | n.s. |
| 21 | n.s. |

d) Extended Data Fig. 2e: p-values of cell densities for *P. deceptionensis* (Pdec) in cocultures with *Chlamydomonas* sp. (Csp) compared to Pdec monocultures in an $NH_4^+$-supplemented YBCII medium.

| Day | Pdec+Csp |
|---|---|
| 0 | n.s. |
| 1 | n.s. |
| 2 | n.s. |
| 3 | ** |
| 4 | ** |
| 5 | n.s. |
| 6 | n.s. |
| 7 | n.s. |
| 11 | n.s. |
| 14 | * |
| 18 | ** |
| 21 | *** |

e) Extended Data Fig. 3b: p-values of cell densities for *Chlamydomonas* sp. (Csp) after one week in different fractions of ΔSM_Csp+Msta compared to Csp grown in non-filtered ΔSM_Csp+Msta.

| Day | < 50 kDa | < 3 kDa |
|---|---|---|
| 7 | n.s. | n.s. |

f) Extended Data Fig. 3c: p-values of cell densities for *M. stanieri* (Msta) after one week in different fractions of ΔSM_Csp compared to Msta grown in non-filtered ΔSM_Csp.

| Day | < 50 kDa | < 3 kDa |
|---|---|---|
| 7 | n.s. | n.s. |

g) Extended Data Fig. 4a: p-values of normalised total chlorophyll content for *Chlamydomonas* sp. (Csp) grown in different spent media compared to Csp grown in an

$NH_4^+$-supplemented YBCII medium. SM_Csp+Msta: Cocultures spent media and ΔSM_Csp+Msta: Heat-treated cocultures spent media.

| Day | Csp+SM_Csp+Msta | Csp+ΔSM_Csp+Msta |
|---|---|---|
| 7 | *** | ** |
| 11 | ** | *** |

h) Extended Data Fig. 4b: p-values of normalised carotenoid content for *Chlamydomonas* sp. (Csp) grown in different spent media compared to Csp grown in an $NH_4^+$-supplemented YBCII medium. SM_Csp+Msta: Cocultures spent media and ΔSM_Csp+Msta: Heat-treated cocultures spent media.

| Day | Csp+SM_Csp+Msta | Csp+ΔSM_Csp+Msta |
|---|---|---|
| 7 | * | * |
| 11 | ** | *** |

i) Extended Data Fig. 5b: p-values of normalised total chlorophyll content of *Chlamydomonas* sp. (Csp) cells grown in monoculture in $NH_4^+$-supplemented YBCII medium, in mono-, co- and tripartite cultures in an $NH_4^+$-depleted YBCII medium, compared to Csp+Msta cocultures grown in $NH_4^+$-depleted YBCII medium. Msta: *M. stanieri*., Vdiaz: *V. diazotrophicus*. n.d.: not detected.

| Day | Csp+$NH_4^+$ | Csp | Csp+Msta+$NH_4^+$ | Csp+Vdiaz | Csp+Msta+Vdiaz |
|---|---|---|---|---|---|
| 7 | * | n.d. | ** | * | * |
| 11 | * | n.d. | ** | ** | * |
| 14 | * | n.d. | *** | ** | ** |
| 18 | ** | n.d. | ** | * | *** |
| 21 | ** | n.d. | **** | * | *** |

j) Extended Data Fig. 5c: p-values of normalised carotenoid content of *Chlamydomonas* sp. (Csp) cells grown in monoculture in $NH_4^+$-supplemented YBCII medium, or in mono-, co- and tripartite cultures in an $NH_4^+$-depleted YBCII medium, compared to Csp+Msta cocultures grown in $NH_4^+$-depleted YBCII medium. Msta: *M. stanieri*., Vdiaz: *V. diazotrophicus*.n.d.: not detected.

| Day | Csp+$NH_4^+$ | Csp | Csp+Msta+$NH_4^+$ | Csp+Vdiaz | Csp+Msta+Vdiaz |
|---|---|---|---|---|---|
| 7 | n.s. | n.d. | ** | ** | ** |
| 11 | * | n.d. | * | * | ** |
| 14 | * | n.d. | ** | ** | ** |

| | | | | | |
|---|---|---|---|---|---|
| 18 | * | n.d. | *** | * | ** |
| 21 | ** | n.d. | ** | ** | ** |

k) Extended Data Fig. 5d: p-values of cell densities for *V. diazotrophicus* (Vdiaz) in cocultures with *M. stanieri* (Msta) compared to Vdiaz in monocultures in an $NH_4^+$-depleted YBCII medium.

| Day | Vdiaz+Msta |
|---|---|
| 0 | n.s. |
| 1 | n.s. |
| 2 | n.s. |
| 3 | * |
| 4 | n.s. |
| 5 | * |
| 6 | n.s. |
| 7 | n.s. |
| 11 | n.s. |
| 14 | n.s. |
| 18 | n.s. |
| 21 | n.s. |
